# Emergence of stable sensory and dynamic temporal representations in the hippocampus during working memory

**DOI:** 10.1101/474510

**Authors:** Jiannis Taxidis, Eftychios Pnevmatikakis, Apoorva L. Mylavarapu, Jagmeet S. Arora, Kian D. Samadian, Emily A. Hoffberg, Peyman Golshani

## Abstract

Hippocampal networks form maps of experience through spiking sequences that encode sensory cues, space or time. But whether distinct rules govern the emergence, stability and plasticity of externally driven and internally-generated representations remains unclear. Using two-photon calcium imaging, we recorded CA1 pyramidal populations across multiple days, while mice learned and performed an olfactory, delayed, working-memory task. We observed anatomically intermixed spiking sequences, comprised of ‘odor-cells’ encoding olfactory cues, followed by ‘time-cells’ encoding odor-specific delay time-points. Odor-cells were reliably activated across trials and retained stable fields over days and different delays. In contrast, time-cells exhibited sparse, unreliable activation and labile fields that remapped over days and extended delays. Moreover, the number of odor-cells remained stable, whereas time-cells increased over days during learning of the task, but not during passive exposure. Therefore, multi-modal representations with distinct learning-related dynamics and stability can co-exist in CA1, likely driven by different neurophysiological and plasticity mechanisms.

## INTRODUCTION

The hippocampus forms representations of the external world, as well as internal representations of planned actions and elapsed time^1-3^. Cell assemblies encoding such information generate spiking sequences which temporally organize experiences and form maps of related memories^4–5^.

An example of internally-generated representations, apart from the widely studied place-cells, is representations of time. These are prominent in delayed-response tasks involving working memory i.e. the active retention of information over a short period. Neuronal ensembles in area CA1 form spiking sequences of ‘time cells’, encoding elapsed time during a delay period^6–10^ which can be specific to the remembered stimulus or the ensuing response^11,12^. Similar sequences have also been observed in area CA3^13^ and cortical regions^14,15^ and may integrate elements of temporally related memories^16^.

Moreover, hippocampal neurons produce selective responses to external sensory stimuli, including visual cues^17^, auditory^18^, olfactory cues^19^ or combinations of those^20^. Olfactory representations in particular (i.e. cells that respond to specific odors) are thought to be involved in various learning processes^19,21–23^. Hippocampal networks receive olfactory information through the lateral entorhinal cortex^24,25^ and project back to olfactory cortex^26^, rendering them a crucial element in odor-discrimination^19,27^, odor-rule learning^28^ or odor-sequence memory^29^.

The principles underlying the emergence and stability of such diverse representations remain largely unknown. On one hand, stable network dynamics (e.g. a fixed set of neurons encoding a specific stimulus) can be robust to small perturbations, allowing information to be reliably conveyed downstream^30^. On the other hand, network flexibility in generating population dynamics (e.g. highly dynamic spiking patterns) allows for a larger pool of potential sequences and quick adaptation to perturbations, and thus a larger amount of information to be encoded^31,32^. Can both of these regimes coexist in the hippocampus or is there a common rule governing the formation, maintenance, and plasticity of network dynamics? For example, do both external (sensory) and internal (temporal) representations activate the same neuronal population and with the same reliability? Do they share common long-timescale stability characteristics or short-term adaptability to contextual changes? Do they both pre-exist in the network or do they emerge as a context is learned? Disentangling these alternatives will yield valuable insights on whether the hippocampus employs a common internal mechanism for multi-modal encoding, or whether it allows for multiple representational regimes, likely driven by distinct neurophysiological mechanisms and synaptic pathways.

Addressing these questions requires studying multi-modal hippocampal representations and comparing their characteristics over long time-scales. To that end, we performed two-photon calcium imaging *in vivo* in the dorsal CA1 pyramidal layer of head-fixed mice^33^ while they performed an olfactory delayed non-match-to-sample task (DNMS) requiring working memory^34^. We recorded activity from thousands of neurons over multiple days during and after learning and across delay periods. We observed two anatomically intermixed groups of pyramidal cells: (i) neurons that were active during the presentation of specific odor stimuli (‘odor-cells’) and (ii) neurons that were active during specific time points in the delay following given odor stimuli (‘time-cells’). Together they formed odor-specific spiking sequences that tiled the odor cue and delay period and encoded stimulus identity and time. Odor-cells exhibited robust activation across trials and retained stable spiking fields for multiple days and delay durations. In contrast, time-cell sequences were less reliable, sparse and dynamic, as time-cells remapped their activity over days and delays. Interestingly, during learning of the task, the number of time-cells increased, whereas the number of odor-cells remained stable. This increase was not observed in untrained animals, where no memory context was present.

Collectively, our findings describe two distinct representational regimes, coexisting spatially intermixed within the same neural network in CA1; a robust and stable, stimulus-driven, sensory code combined with a sparse and dynamic, memory-specific temporal code. The former pre-exists exposure to task context while the latter mostly emerges during learning. Therefore, hippocampal networks can exhibit combined stable externally-driven and flexible internally-driven representations, suggesting that these likely emerge through separate synaptic pathways and distinct neurophysiological mechanisms.

## RESULTS

### Two-photon calcium imaging in dorsal CA1 during an olfactory working-memory task

Adult (11-31 weeks old) Gad2Cre:Ai9 and Gad2Cre:Ai14 mice (N = 5 and 3 respectively), expressing tdTomato in GABAergic interneurons, were injected with AAV1-Syn-GCAMP6f virus in the right dorsal CA1 area (dCA1). A titanium ring, with a glass coverslip at the bottom, was implanted over the corpus callosum after surgical aspiration of the overlying parietal cortical region (Figure 1A). After recovery, mice were water-deprived and trained to perform an olfactory delayed non-match-to-sample task (DNMS) involving working memory^34^. Mice were head-fixed on a spherical treadmill (Figure 1A) where a lickport system delivered odorized and clean air as well as water droplets. Two odor cues were presented for 1 sec each, with a 5 sec delay period in between (Figure 1B). Mice were trained to lick the lickport for water reward if the two odors did not match and refrain from licking when they matched. Lick responses were assessed during a 2 sec response window, 1 sec after the second odor presentation. Two odors were used throughout: isoamyl acetate (‘odor-A’) and pinene (‘odor-B’). Each behavioral session was comprised of 20 trials with an equal number of all four odor combinations, randomly distributed. Performance was assessed as the ratio of summed correct hits and correct rejections over all trials (Figure 1C).

**Figure 1:**
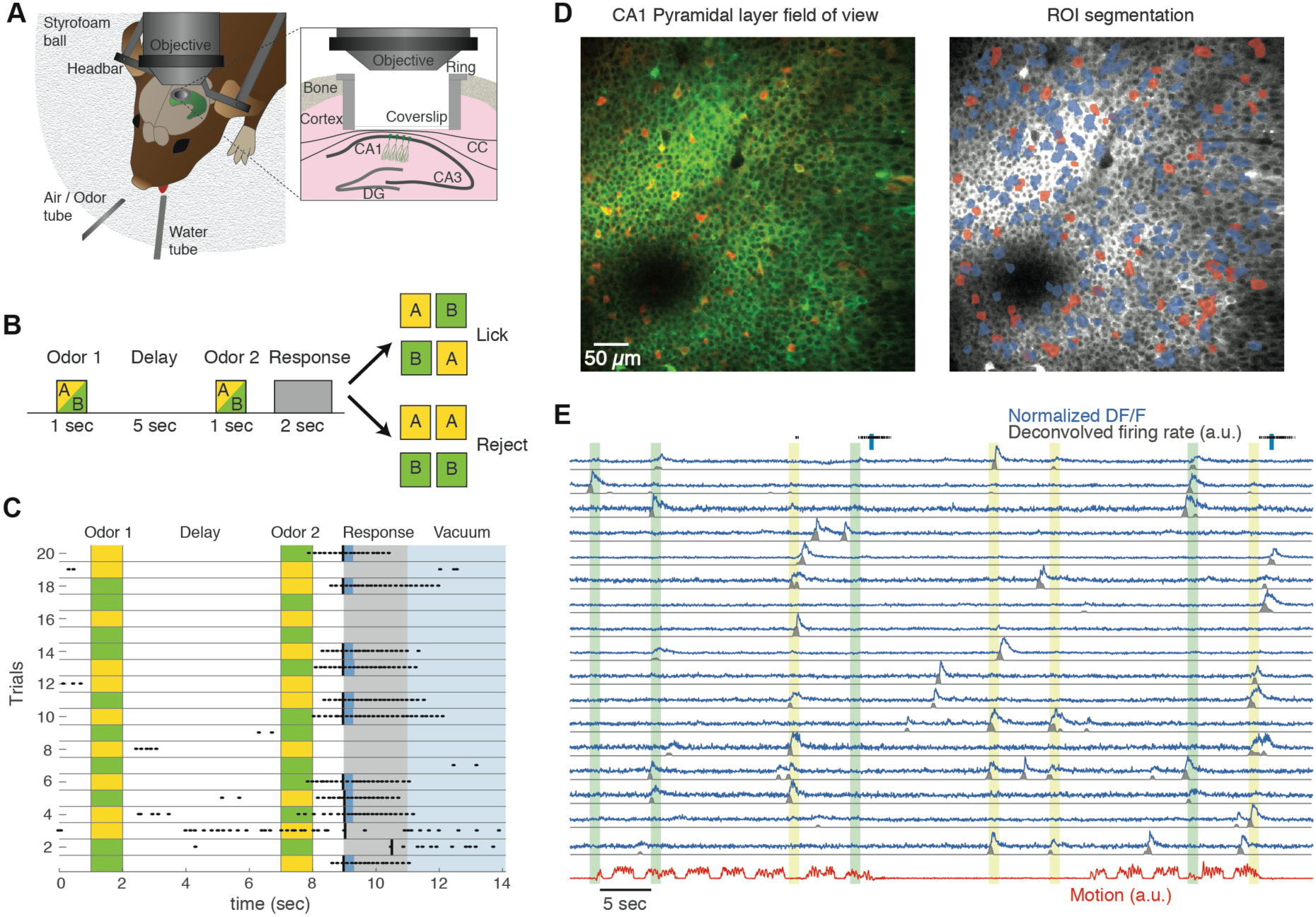
Experimental setup and recordings. **A.** Illustration of the behavioral and experimental set-up. The mouse is head-fixed through a headbar on a spherical treadmill under the two-photon microscope objective. A tube delivering clean or odorized air and another delivering water rewards is placed in front of the mouse. A third tube (not shown) applies vacuum at the end of the trial to clear remaining water drops and odorized air. A titanium ring with an attached coverslip is placed over the dorsal hippocampus, after surgical aspiration of the overlying cortex (inset), allowing imaging of the dorsal CA1 pyramidal layer. CC: Corpus callosum. **B.** Schematic of a behavioral trial. Two odors are presented for 1 sec each, with a 5 sec delay between them. Each odor can be either isoamyl acetate (‘Odor-A’, yellow) or pinene (‘Odor-B’, green). The animal’s response is assessed during a 2 sec window after the second odor. If the two odors do not match, it can lick the water tube to release a water reward (‘hit’). If the odors match it needs to withhold licking (‘correct rejection’). **C.** Example session comprising of 20 trials from one animal. Odor sequences indicated by yellow and green colors and licks by black dots. Odors are presented in random order. Animals rarely lick during the delay period. The first lick during the response window in non-match trials (black line) will trigger a water drop (blue) in non-match trials. Licking during match trials does not trigger a reward (‘false alarm’). Each trial ends with a 3 sec application of vacuum to clear the water tube and lingering odors. **D.** Left: Example template (average) frame from a motion-corrected field of view of dCA1 pyramidal layer, recorded in green and red channels, from one Gad2Cre:Ai9 mouse expressing GCaMP6f (green) after viral transfection with AAV-Syn-GCAMP6f virus. GABAergic interneurons express Td-Tomato (red). Right: Same field of view after segmentation of ROI with the CNMF algorithm (Methods). Blue: pyramidal cells that produced calcium transients which were detected by the CNMF algorithm. Red: GABAergic cells detected from their mean (non-activity-based) fluorescence in the red PMT channel (Methods). **E.** Example normalized ΔF/F traces (blue) from pyramidal cells during 4 trials in a recording session. The binned deconvolved spiking probability (‘firing rate’) of each trace is shown in black and simultaneously recorded motion on the ball in red. Licking (black marks) and water delivery (blue bars) are depicted on top.

We performed resonant scanning *in vivo* two-photon calcium on the dCA1 pyramidal layer, recording activity from hundreds of cells per mouse, while animals performed the DNMS task (Figure 1D). Calcium imaging movies were recorded on green and red PMT-channels to segregate pyramidal and interneuronal activity (Figure 1D). Using custom-written software, based on an extended version of the CNMF algorithm^35,36^, movies were motion-corrected, active regions of interest (ROI) were segmented and split into pyramidal cells and interneurons. Calcium signals were extracted and deconvolved to yield a final binned spiking probability measure (Figure 1E). We use this time-binned measure as a proxy of spiking activity and refer to it as ‘firing rate’ for simplicity throughout the text. On average, each recording session (74 sessions in total, typically consisting of 100 trials) yielded 242.1 ± 137.6 pyramidal cells (mean ± SD) and 47.1 ± 14.9 interneurons (79.75% ± 10.8% and 20.25% ± 10.8% of all ROI respectively). Only 2.2% ± 2% of all CNMF-detected ROI were interneurons, as calcium traces from these cells did not meet CNMF criteria (Methods). All interneurons were removed from our ensuing analysis; only pyramidal cell activity was analyzed further. Furthermore, the same field of view was recorded during every imaging session and was registered across days, tracking the activity of the same neurons for up to 11 consecutive days. Finally, locomotion on the treadmill was also monitored (Figure 1E).

To ensure that mice relied on working memory to perform the task we conducted a series of control measurements and experiments. First, mice did not participate when the odors where turned off, excluding the use of auditory cues from air valves. To ensure mice did not rely on a comparison of airflow values, those were set to be the same for the two odors and also performance was not affected by randomly altering them for the two odors (Figure S1). To ensure mice did not detect odor mixture during the second odor presentation in non-match trials, we minimized the common path for the two odors (~4cm) and used a photo-ionization detector (PID) to confirm that odorant from the first stimulus was not present in the second one (Figure S1). Next, to ensure mice did not detect lingering odor during the delay, they were trained on a non-match-to-long-duration-sample task^34^ (N = 3) where a long 5 sec odor-stimulus was followed by a 1 sec matching or non-matching odor and mice had to lick for reward only in non-match trials (Methods). Mean performance dropped to chance levels when the odorant concentration of the long stimulus was reduced to 0.01%, yielding a threshold for odor detection during the DNMS delay period. Based on this threshold, odorants were undetectable by mice within 1 sec of odor offset in DNMS (Figure S1). Finally, performance declined when increasing the delay duration (N = 4 mice), corroborating that working memory was employed to perform the task (Figure S1).

### Pyramidal cells form spiking sequences encoding odor identity and odor-specific delay time

We first focused on sessions where mice performed at well-trained levels, (average performance per day ≥90%; 32 imaging sessions total) and, in each trial, we examined activity between the first odor cue onset and the delay offset, i.e. cue presentation and working-memory intervals respectively. We observed ‘odor-cells’ that fired during the presentation of a specific odor (Figure 2A), and odor-specific ‘time-cells’ that fired during specific time bins in the delay after a given odor (Figure 2B). Collectively, these groups formed two distinct sequences depending on the initial odor (‘trial-type’): ‘A-sequence’, comprised of ‘odor-A’ and ‘time-A’ cells, spiking during the presentation of odor-A and its ensuing delay, and ‘B-sequence’ correspondingly for odor-B and its ensuing delay. Cells in each sequence (‘sequence-cells’) remained overall silent during the opposite trial-type (Figure 2C) and were triggered by the second odor as well, though behavioral and reward variations confounded that activity (Figure 2C). With our classification criteria, 22.6% ± 7% of detected ROI per session were classified as sequence-cells. 5% ± 2% ROI were odor-A cells, 8.3% ± 3.7% were time-A, 5.5% ± 2.2% were odor-B and 5.3% ± 2.8% were time-B (Figure 2D). Finally, 93.8% ± 6.1% sequence-cells per session had a single preferred trial-type, while 6.2 % ± 6.1% had a significant field in both sequences.

**Figure 2:**
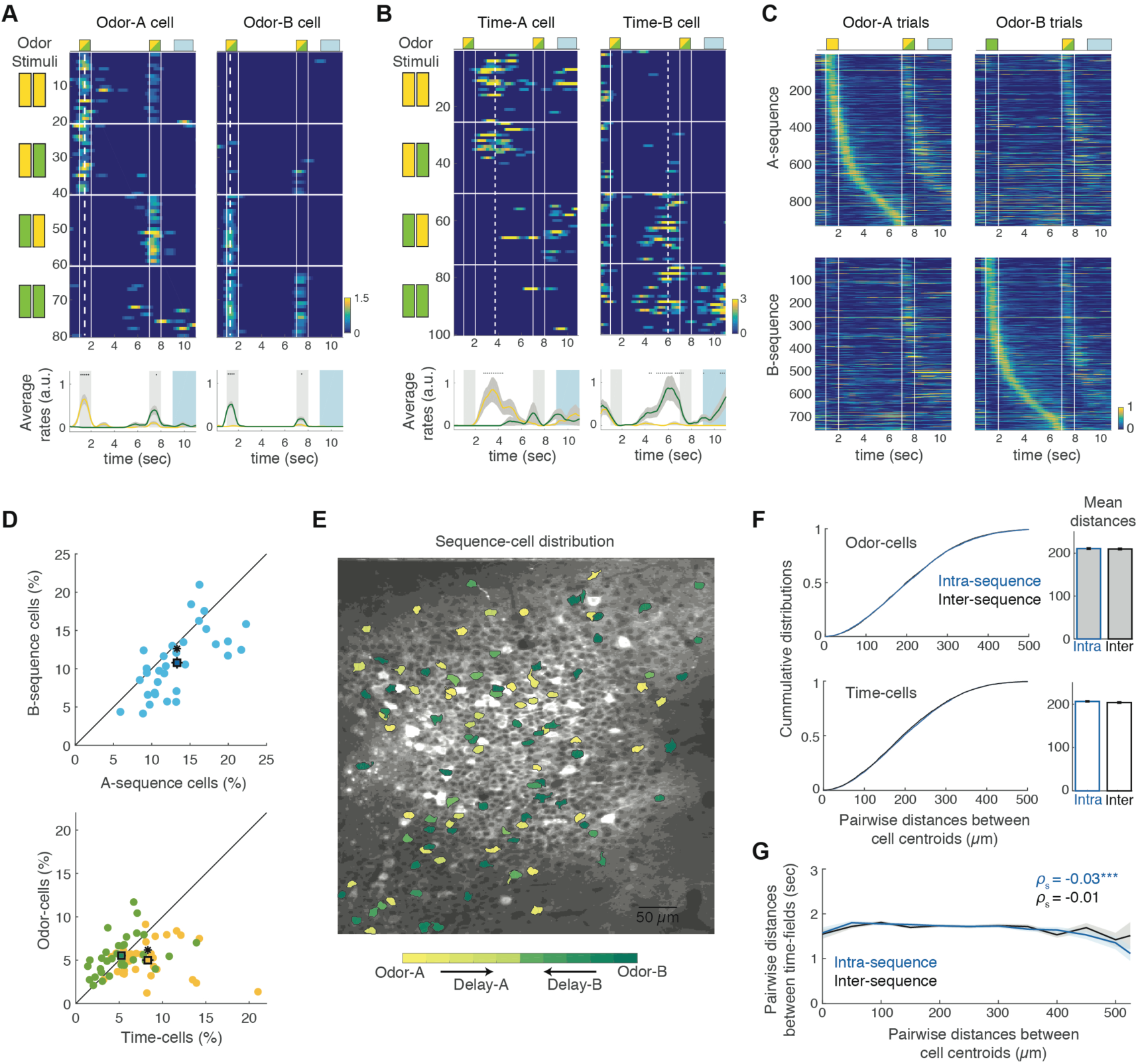
Uniformly distributed dCA1 pyramidal cells form spiking sequences encoding odor and odor-specific delay time. **A.** Two example ‘odor-cells’ encoding the presentation of odor-A (isoamyl acetate; Left) or odor-B (pinene; Right). Each row represents the neuron’s estimated firing rate during a trial. Trials are stacked in four blocks (horizontal lines) according to the odor combination, shown in left (odor-A in yellow; odor-B in green). Vertical lines: two odor delivery blocks followed by the response window (trial layout shown on top). Dashed lines: firing field (location of maximum average firing rate over all preferred trials). Firing rates are normalized by the neuron’s average rate at its field. Bottom: Mean ± SE firing rate of the cell during all odor-A trials (yellow) and odor-B trials (green). Dots: time bins with significant difference between the two mean rates (P < 0.05, Wilcoxon test (WT); FDR). Grey bars: odor delivery. Blue bars: response window. **B.** Same as **A** for two example ‘time-cells’ encoding time points during the delay following a specific odor. **C.** Top: Mean normalized firing rates from A-sequence and B-sequence-cells pooled from all recording sessions, during odor-A trials (left) and odor-B trials (right). **D.** Top: Ratio of A-sequence versus B-sequence-cells per imaging session. A-sequence comprised of a higher percentage of cells on average (rectangle: mean ± SE; P < 0.05, paired-sample t-test). Bottom: Ratio of odor-A versus delay-A cells (yellow) and odor-B versus delay-B cells (green). Asterisk: P < 0.01, paired-sample t-test, Bonferroni corrected over the two tests. **E.** Example field of view from one recording session, where all the sequence-cells are outlined, superimposed on the mean template, color coded by their firing field. **F.** Cumulative distributions of pairwise distances between all odor-cell centroids (top) or time-cells (bottom) belonging to the same sequence (blue) or opposite sequences (black). Inset: Distribution means ± SE. No significant differences were found (P > 0.05; Cumulative distributions: Two-sample Kolmogorov-Smirnov test; Means: WT; FDR). **G.** Mean ± SE of pairwise (binned) time-distance between the fields of time-cells of the same sequence (blue), and between time-cells of opposite sequences (black) as a function of their centroid distance. *ρ*s: Spearman correlation, asterisks: P < 0.001, permutation distribution test.

To ensure sequence-cells did not reflect any lingering odorant concentration declining during the delay, we recorded from sessions with either the default 5% odorant concentration, 1% or 0.5% concentrations (N = 3 mice). If each sequence-cell encoded a particular concentration value, then by reducing the odorant concentration, odor-cells would stop spiking and time-cell activity would be shifted forward in time (as the corresponding lower concentration would occur earlier in the delay). However, we found that sequences remained robust with cells retaining their firing field and mean firing rate during all concentrations (Figure S2), indicating that activity was independent of odorant concentration.

We next asked whether the sequences encoded locomotion on the ball. Individual animals exhibited different locomotion patterns during trials (though all mice yielded odor- and time-cells), with some initiating running at the first odor and throughout the delay and others halting during the odors, suggesting a dissociation between motion and sequence spiking. Indeed, when ranking trials based on the total locomotion of a mouse (all N = 8 mice) and comparing sequences during the 30% lowest- versus 30% highest-locomotion trials, we found no difference in spiking (Figure S3), suggesting that both olfactory and temporal representations were not confounded by locomotion.

Finally, we tested whether these sequences were triggered by auditory stimuli related to air valves switching between clean and odorized air. We recorded activity after turning the odorized air supply off, while retaining all valve auditory cues (N = 3 mice). Sequence-cells from preceding sessions became overall silent when the odors were turned off (P < 0.05; Wilcoxon test (WT), FDR), indicating that their activity does not encode auditory cues (Figure S4).

Collectively, these results indicate that odor-cells encode the identity of an olfactory cue whereas time-cells encode time bins during the delay after a given odor, thus effectively carrying information about time and stimulus identity held in working memory. Are the two representations part of a continuous sequence, triggered by odor-cells? Or are they generated by district neurophysiological mechanisms? We address this question in the following sections.

### Odor and time-cells are uniformly distributed over the dCA1 pyramidal layer

Place fields are uniformly distributed along the CA1 pyramidal layer^33^, but the anatomical distribution of sensory- and time-encoding remains unexplored.

Sequence-cells were typically intermixed and distributed throughout the pyramidal layer in our field of view (Figure 2E). To test whether odor-cells lying nearby are more likely to encode the same odor versus opposite ones, we compared the centroid distances of all odor-cell pairs in the same sequence (intra-sequence), with distances from all odor-cell pairs of opposite sequence (inter-sequence). The same analysis was performed for time-cells (Figure 2F). We found no significant differences in the distance distributions for either category (P > 0.5, Kolmogorov-Smirnov test for cumulative distributions and WT for means; FDR), indicating that proximity between two cells was not related to their preferred odor.

We next examined whether time-cells lying nearby had a higher probability of having closer time-fields. We computed time-distances between the fields of each time-cell pair of the same sequence (intra-sequence), as a function of their centroid distance. The same analysis was performed for cells belonging to opposite sequences (inter-sequence). The inter-sequence distribution was uniform (P > 0.05) whereas intra-sequence field-distances exhibited a very small but significant rank correlation with centroid-distances (P < 0.001; Spearman (rank) correlation; permutation distribution test; FDR; Figure 2G).

Our results suggest that sequence-cells are overall uniformly distributed along the dCA1 pyramidal layer, with no spatial clustering involved in encoding properties of cells.

### Robust odor-representation is followed by degrading time/odor-encoding during the delay

Do both odor- and time-representations exhibit common spiking characteristics and decoding power? We sought to compare the information carried by odor and time-cells individually and on the ensemble level.

First, olfactory representation was redundant whereas time-representation was sparse. Given that a comparable number of cells was on average classified as odor- or delay-encoding, a larger number of cells had their firing field at each of the odor time bins compared to each delay time bin (Figure 3A). Interestingly, the number of firing fields decreased as a function of delay time and could be fitted by a power law distribution with a = −2.68 exponent (P < 0.01 for goodness of fit based on random sampling; Methods).

**Figure 3:**
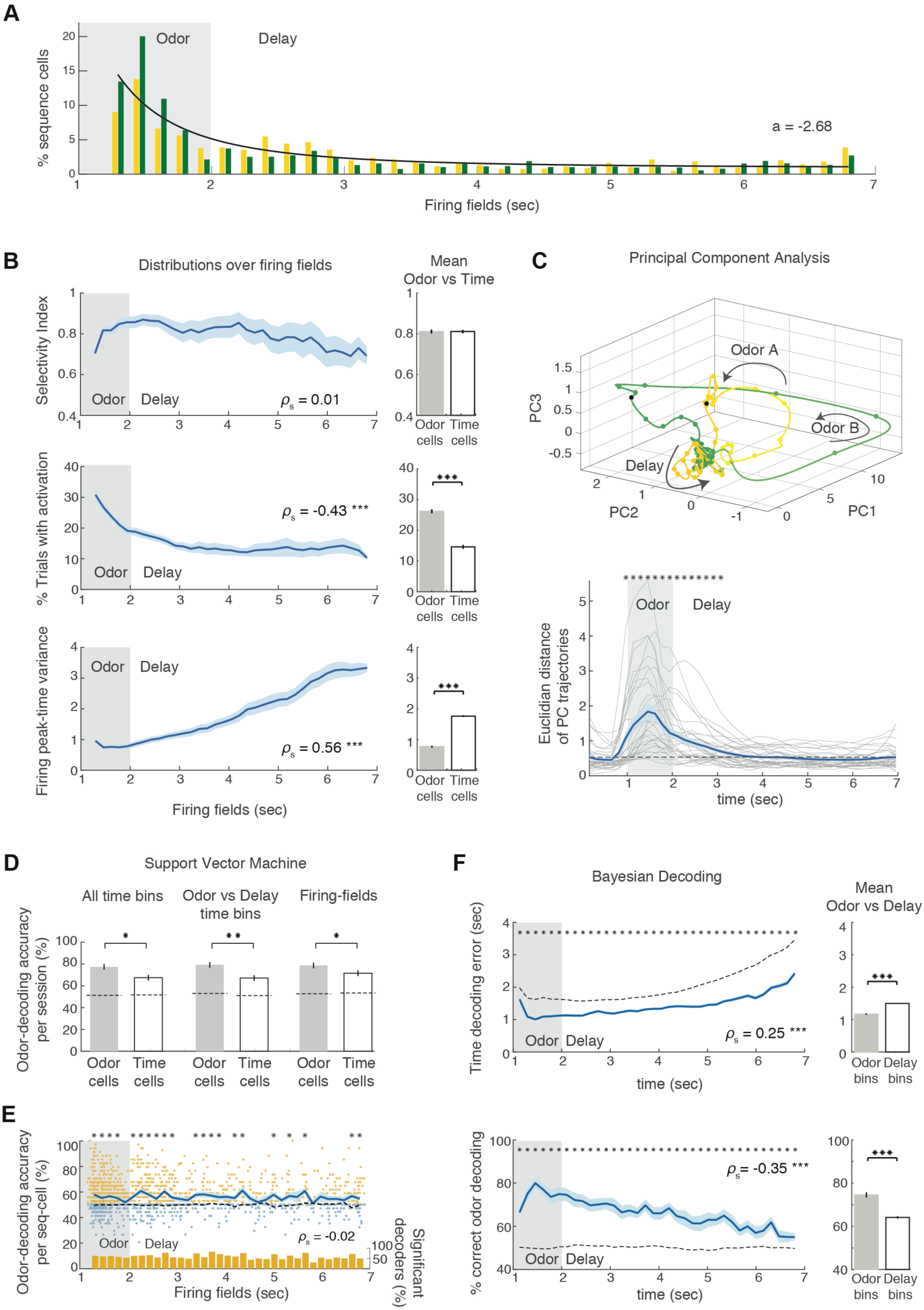
Robust odor-cell activity is combined with sparse time-cell spiking and progressive information loss during the delay. **A.** Distribution of firing fields for the two sequences (yellow and green respectively) and power-law fit (black) of mean distribution with exponent ‘a’ shown in inset. **B.** Mean ± SE distributions of selectivity index (top), percentage of preferred trials where each cell spiked within its field (middle) and variance of each cell’s firing peak-time per trial (bottom) as a function of the cell’s field. All curves were smoothed with a 5-point moving average. Right: Mean ± SE of each corresponding measure over all odor- and time-cells. Asterisks: P < 0.001; WT; FDR. **C.** Top: Mean PCA trajectory along the first 3 PCs, during all A-trials (yellow) and B-trials (green) from one session, spline interpolated over time bins (dots). Darker colors indicate later time bins. Arrows indicate time direction. Black dots signify delay onset points. Bottom: Mean Euclidean distance between the average PCA trajectories of each session (grey) and over all sessions (blue; mean ± SE). Asterisks: Time bins where the mean distance is significantly higher than baseline (dashed line P < 0.05; WT; FDR). **D.** Odor-prediction accuracy by SVM classifiers using all odor- or all time-cell firing rates of each session, averaged over the odor-delay interval (left), or just the odor or delay time bins respectively (middle), or by using only each cell’s field time bin (right). In each case, the prediction accuracy of each group is significantly higher than chance baseline (dashed lines; P < 0.001, WT; FDR). Collective odor- or time-cell activity yields significantly higher prediction accuracies (Asterisks: * P < 0.05, ** P < 0.01, paired sample t-test for first comparison and WT for the rest; FDR). No significant differences were found between predictions from the same cell group over the three different time intervals (P > 0.05; WT; FDR). **E.** SVM odor-prediction accuracy using single sequence-cell firing rates over the odor-delay interval, plotted as function of each cell’s field. Cells with significantly higher prediction accuracy than baseline are shown in yellow (P < 0.05; tailed WT, FDR), and those with non-significant one in blue (small random noise is added to their field location for clarity of overlapping data points). Blue curve: mean ± SE for cells of each firing field shows no increasing or decreasing trends. Asterisks: significant mean prediction accuracy against mean baseline (black; P < 0.05; tailed paired sample t-test or WT depending on distributions – see Methods; FDR). Histogram depicts the ratio of significant decoder cells in each field (*ρ*s: Spearman correlation). **F.** Mean ± SE Bayesian time-decoding error (top) and odor-decoding accuracy (bottom) using the firing rates of all sequence-cells in each session, as a function of time. Asterisks: At each time bin, the average time prediction error is significantly lower and the odor-decoding accuracy significantly higher than their corresponding chance baselines (dashed curves; P < 0.05; WT; FDR). Time errors increase monotonically and decoding accuracy drops monotonically (*ρ*s: Spearman correlation; Asterisks: P < 0.001; permutation distribution test). Right: Mean ± SE of each measure over all odor- and time-bins (asterisks: P < 0.001, WT; FDR).

To quantify the firing field properties of individual sequence-cells, we computed their selectivity index (SI), their activation probability during their preferred trials and their spiking time variance as a function of their firing field (Methods). The mean SI did not exhibit a significant trend over firing fields (P > 0.05, permutation distribution test for Spearman correlation) indicating a similar selectivity to their preferred trials for all cells. But, the activation probability monotonically decreased and the variance increased as a function of firing field time (P < 0.001; FDR), so that odor-cells had significantly higher activation probability and lower spiking variance that time-cells on average (P < 0.001; WT; FDR; Figure 3B).

These findings suggest a potential loss of collective information by sequences along the delay period. Indeed, when applying principal component (PC) analysis to assess the PC-space trajectories for the two trial types (see Methods), the average trajectories of the two trial-types deviated during odor presentation but converged and become intermingled as delay progressed (Figure 3C). Consequently, during odor presentation and early delay there was a significant average Euclidean distance between the mean A-trial and B-trial trajectories on the full PC space (P < 0.05), beyond which it dropped to chance-levels (P > 0.05; WT; FDR; Figure 3C).

Individual PC trajectories were affected by sparse time-cell activation and the variability in activation patterns of time-cells during individual trials, leading to the converging mean trajectories. But could odor information be efficiently retained within the ensemble activity throughout the delay? To decode odor identity from either odor-cells or time-cells collectively, we implemented a binary support vector machine (SVM) classifier. We first used firing rates from each group, averaged over the entire odor-delay interval, to train the classifier. We then narrowed our analysis by retraining the classifier using activity over the odor-presentation and delay period respectively, and finally using only the field time-bin of each cell (Figure 3D). Odor-decoding accuracy was significantly above chance for both cell groups using all three methods (P < 0.001 WT versus shuffled data; FDR), demonstrating that odor-information can be efficiently extracted from either cell group. Odor-decoding accuracy with each cell group was comparable for all three methods (P > 0.05) but, intuitively, odor-cells always yielded significantly better decoding than time-cells (P < 0.05; WT; FDR).

Can odor information by efficiently decoded from individual cell activity as well? When applying an SVM classifier on the firing rate of each sequence-cell separately over the entire odor-delay interval (Figure 3E), we found that the ratio of classifiers performing significantly better than chance and the mean SVM accuracy in each firing field were uniform over the delay interval (Spearman correlation: *ρ*s= −0.02; P > 0.05 random permutation test), reflecting the selectivity uniformity over all fields. However, the firing fields that yielded significant mean accuracy over all local classifiers (P < 0.05 vs chance baseline; WT, FDR), were sparser in later segments of the delay than earlier ones or during the odor-presentation.

Since SVM classification relies solely on the activation of cells over a given time interval but ignores the timing of this activation, we used Bayesian decoding to extract information on time by the ensemble activity. Each decoder was trained over the mean activity of all sequence-cells, concatenated over A-trials and B-trials and decoded time bins along twice the odor-delay interval, representing time as well as odor-information (see Methods). The mean decoded-time errors (absolute values) were significantly lower and the decoded-odor accuracy significantly higher than their respective chance baselines for all time bins (P < 0.05 WT over shuffled data; FDR), indicating that both types of information were efficiently retained in the collective activity throughout the trial. However, the time-error monotonically increased and the odor-accuracy monotonically decreased over time (P < 0.001, Spearman correlation random permutation; FDR), suggesting a gradual loss of decoding power along the delay.

Collectively, these findings reveal a redundant and reliable code for odor-stimuli yielding high decoding power, and a sparse, more variable temporal code, with odor/time information degrading over the delay period, leading to progressive loss of decoding power.

### Odor-cells remain stable across multiple days whereas time-cells shift their fields

CA1 place and contextual representations are typically dynamic across days, drifting to different neural populations^37–39^. We asked whether olfactory and temporal representations exhibit similar drifts over long timescales of multiple days.

To address this question, we imaged the same field of view for up to 8 consecutive days (29 imaging sessions total) while mice performed the DNMS task (N = 6). The number of active ROI remained overall stable across days and a similar percentage of those were classified as sequence-cells daily (Figure S5), suggesting that activity in the imaged region did not change or deteriorate over multiple imaging sessions. Mean templates from each pair of days were aligned, their corresponding ROI were registered (Figure 4A; Methods) and recurring sequence-cells were detected (i.e. cells belonging to the same sequence both days). The number of matched ROI and the ratio of recurring sequence-cells decreased with longer distance between two days (Figure S5; P < 0.05 and 0.001 respectively; permutation distribution test), suggesting a progressive drift in ensemble activity over time, supporting previous studies^39^.

**Figure 4:**
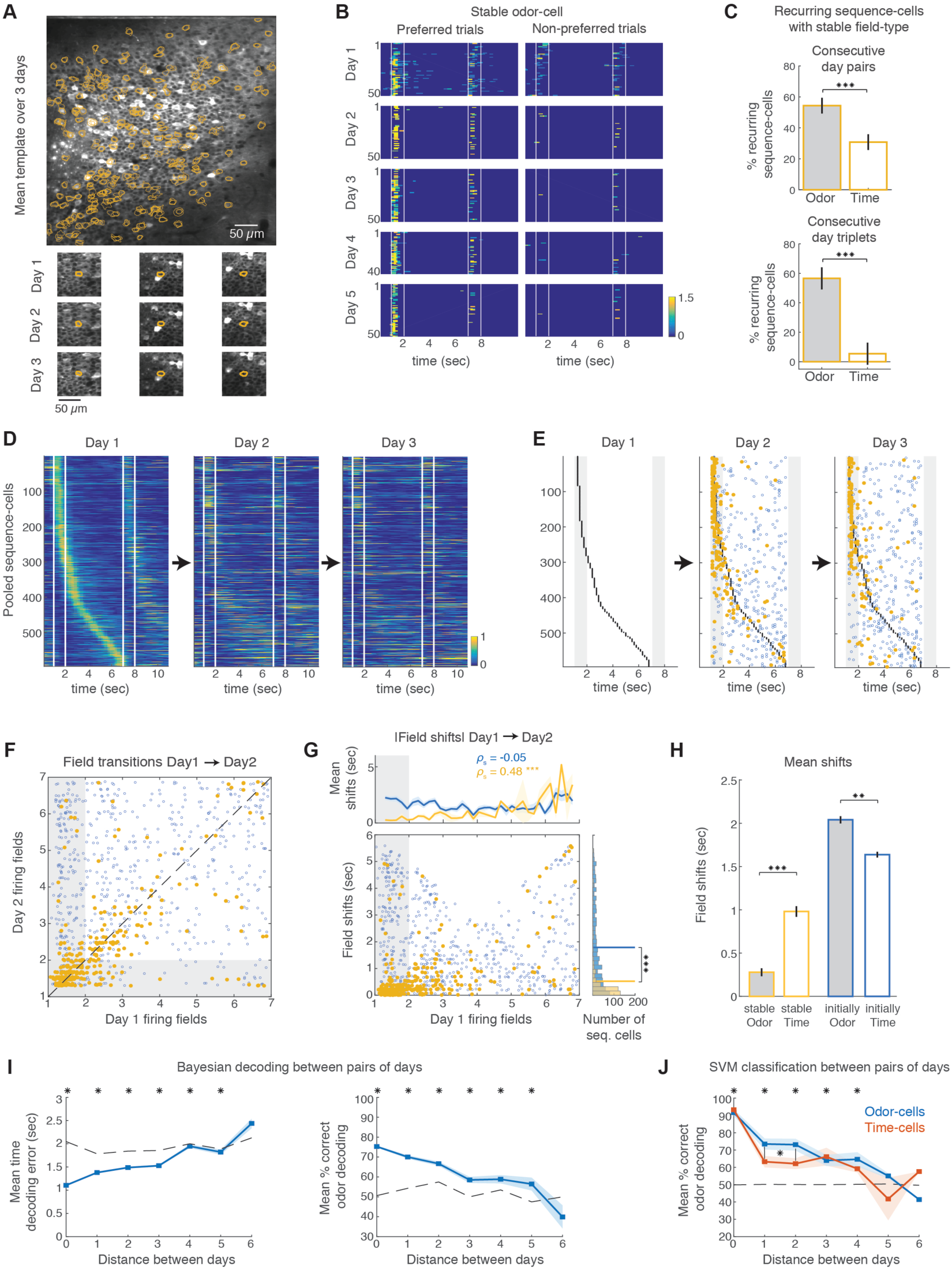
Odor-cells retain stable fields across multiple days whereas time fields are dynamic. **A.** Example template (mean) field of view from one animal, averaged over 3 consecutive days of imaging. Overlapping contours indicate ROI that were detected each day and were registered across days. Bottom: Example registered ROI for each day. **B.** Firing rate over each preferred and non-preferred trial for an example neuron that was classified as an odor-cell for 5 consecutive days. **C.** Top: Mean ± SE percentage of common sequence-cells between two consecutive days that were odor-cells in both days (grey) versus time-cells in both days (white). Bottom: Same for all consecutive day triplets. In both cases, a significantly higher ratio of recurring sequence-cells were odor-cells (P < 0.001, WT and paired sample t-test respectively). **D.** Pooled sequence-cells firing rates from day 1 of all consecutive day triplets, plotted in the same order for the next two days. **E.** Same as **D** but showing only the locations of the original fields (black dots) and the time-bin of their maximum mean firing rate in following days. Blue dots: time-bins from cells that did not retain a significant field on the following day. Yellow dots: time-bins from cells with significant fields on the following days as well (yellow). **F.** Time location of each cell’s mean firing rate peak on Day X as a function of its initial field on Day X+1. Colors as before. Dashed line: Diagonal, indicating no change in firing field. For plotting clarity, a small gaussian noise was added to each data point to avoid overlaps. **G.** Time shifts (absolute values) of sequence-cells from Day X to Day X+1 as a function of their Day X field. Top: Mean ± SE shifts as a function of sequence-cells’ Day X field. Colors as before. *ρ*s: Spearman correlation; asterisks: P < 0.001, permutation distribution. Right: Histogram of field shifts between two days. Lines indicate distribution means. Asterisks: P < 0.001, two-sample Kolmogorov-Smirnov test. **H.** Mean ± SE shifts between Days X - X+1 for odor-cells (grey bars) and time-cells (white bars) for all stable sequence-cells between the two days (yellow; P < 0.001, WT) and initial sequence-cells without significant field the following day (blue outline; P < 0.01, two-sample t-test, Bonferroni over the two tests). **I.** Mean ± SE cross-day Bayesian time-decoding error (left) and odor-decoding accuracy (right) using recurring sequence-cells to train the decoder on Day X and decode Day Y, as a function of distance between days. Distance 0 corresponds to within-day decoding. Asterisks: Significance versus baseline (dashed line), created by 1000 shuffles of the cell identities for all trials of Day Y (P < 0.05; tailed WT; FDR). **J.** Mean ± SE cross-day SVM odor-decoding accuracy using recurring odor-cells or time-cells to train the classifier on Day X and decode Day Y, as a function of distance between days. Asterisks: Significance over baseline (dashed line, identical for both cell groups), created by 1000 shuffles of odor-identities of all trials of Day Y (P < 0.05; tailed WT; FDR). Asterisks correspond to both cell groups. Odor-cell based accuracy is higher than time-cell based for 1 and 2 days apart (asterisk, P < 0.05; WT; FDR. No comparison was performed for 6-days distance since only one data point exists).

Many odor-cells were stable for multiple days, retaining a significant field in their preferred trials (Figure 4B), whereas few time-cells recurred for more that 2 days and almost none for > 3 days. As a result, sequence-cells recurring between two or three consecutive days had higher probability of being odor-cells each day than time-cells (Figure 4C; P < 0.001; WT; FDR). When examining all pairs of days, the number of recurring odor-cells decreased the further apart two days were (P < 0.05) but was higher than that of time-cells, up to 4 days apart (Figure S5).

We examined how this difference in stability of the two cell groups is reflected on ensemble activity. For every triplet of continuous days (‘Days 1-3’), we pooled all sequence-cells in Day 1 that were matched to ROI on the next two days (Figure 4D) and split them into ‘stable’ ones (matching ROI on Days 2 and 3 are still sequence-cells) and ‘unstable’ ones (matching ROI lack a significant field). For the latter group, we used the maximal-firing time-bin over the corresponding trial types to quantify ‘field-shift’ between days (Figure 4E). Again, many Day 1 odor-cells retained their firing field on Days 2 and 3, whereas most time-cells either lost their field on following days or it shifted to a new time bin. To examine the relationship between these shifts and initial field location, we pooled all pairs of consecutive days and again split recurring cells into stable and unstable ones. Again, most stable cells were odor-cells or early time-cells (up to ~1 sec into delay time), whereas most time-cells had shifted their field on Day 2 or lost it (Figure 4F). Consequently, stable-cells exhibited lower shifts on average than unstable ones (Figure 4G; P < 0.001; WT). Moreover, there was a significant relationship between the field bin of a stable cell on Day 1 and its shift on Day 2, with later fields exhibiting higher shifts (Figure 4G; P< 0.001; Spearman correlation), whereas unstable cells exhibited similar shifts irrelevantly of their original field bin. Consequently, stable odor-cells exhibited significantly lower shifts than stable time-cells, whereas unstable odor-cells exhibited higher shifts in than unstable time-cells (Figure 4H), indicating that unstable odor-cells remapped their activity across the entire odor-trial interval (yielding longer shifts).

Finally, we examined how information contained in the ensemble activity evolves across days. First, we performed Bayesian decoding over every pair of days, training the decoder using recurring sequence-cells from day X and decoding their activity on day Y. We found that the further away two days were, the higher the average decoded-time error, and the lower the odor accuracy (Figure 4I). However, both measures remained significantly better than chance up to 5 days apart (P < 0.05, WT on shuffled data; FDR).

Interestingly, even though timing information is progressively lost, only a small minority of recurring sequence-cells tended to transition between odor and time-encoding, and even fewer cells switched preferred trials (Figure S5). To quantify odor-information retained by each cell group across days, we applied an SVM decoder, trained on odor-cell activity during day X and decoding their matched activity on day Y. We repeated the analysis with time-cells separately. Again, decoded-odor accuracy in both groups dropped with increasing distance between days, but remained significantly above baseline up to 4 days apart. Moreover, classifiers trained on odor-cells performed significantly better than those trained on initial time-cells up when decoding activity up to 2 days apart (P < 0.05).

Collectively, our analyses reveal that sensory coding of odor identity is strikingly stable across multiple days, with many odor-cells retaining their field for multiple days. In contrast time-cells were highly dynamic, either shifting their field or losing it. Odor-cell stability combined with a low rate for time-cells of switching preferred odor, odor/time information was retained in ensemble activity even up to multiple days later, albeit with decreasing efficiency.

### Odor-cells remain stable when extending the delay period whereas time-cells shift their fields

To disentangle whether both odor- and time-cells are driven by a common feedforward mechanism, we examined how these sequences adapt to changes in behavioral context over very short timescales of a few minutes. Specifically, if the delay is extended, how will the two representations be affected?

We performed a series of imaging sessions (N = 5 mice; 10 imaging sessions in total) where the delay was extended from 5 to 10 sec within consecutive behavioral sessions. Mice had not been exposed to extended delays previously but quickly adapted (usually within the first 20-trial session) and exhibited a minor but significant reduction in average performance (88.3% ± 14.8%; versus 96% ± 7% for 5 sec trials; P < 0.01; WT). We detected sequence-cells based on the initial 5 sec sessions of each day and examined their activity over the 10 sec ones. Odor-cells generally retained their activity while time-cells tended to shift their spiking throughout the extended time period (Figure 5A). Most of the ‘stable cells’ (i.e. retaining significant field with extended delay) were again odor-cells and early time-cells and exhibited field-shifts that increased as a function of initial field bin, but were on average significantly lower those of ‘unstable’ cells (i.e. without a field in extended delay; P < 0.001; Kolmogorov-Smirnov test). The latter consisted mostly of time-cells with their maximal-firing time-bin shifted forwards or backwards in time, independently of the original field bin (Figure 5B-D). As before, the average shifts of stable odor-cells were again lower than those of stable time-cells whereas the opposite held for unstable ones (Figure 5E).

**Figure 5:**
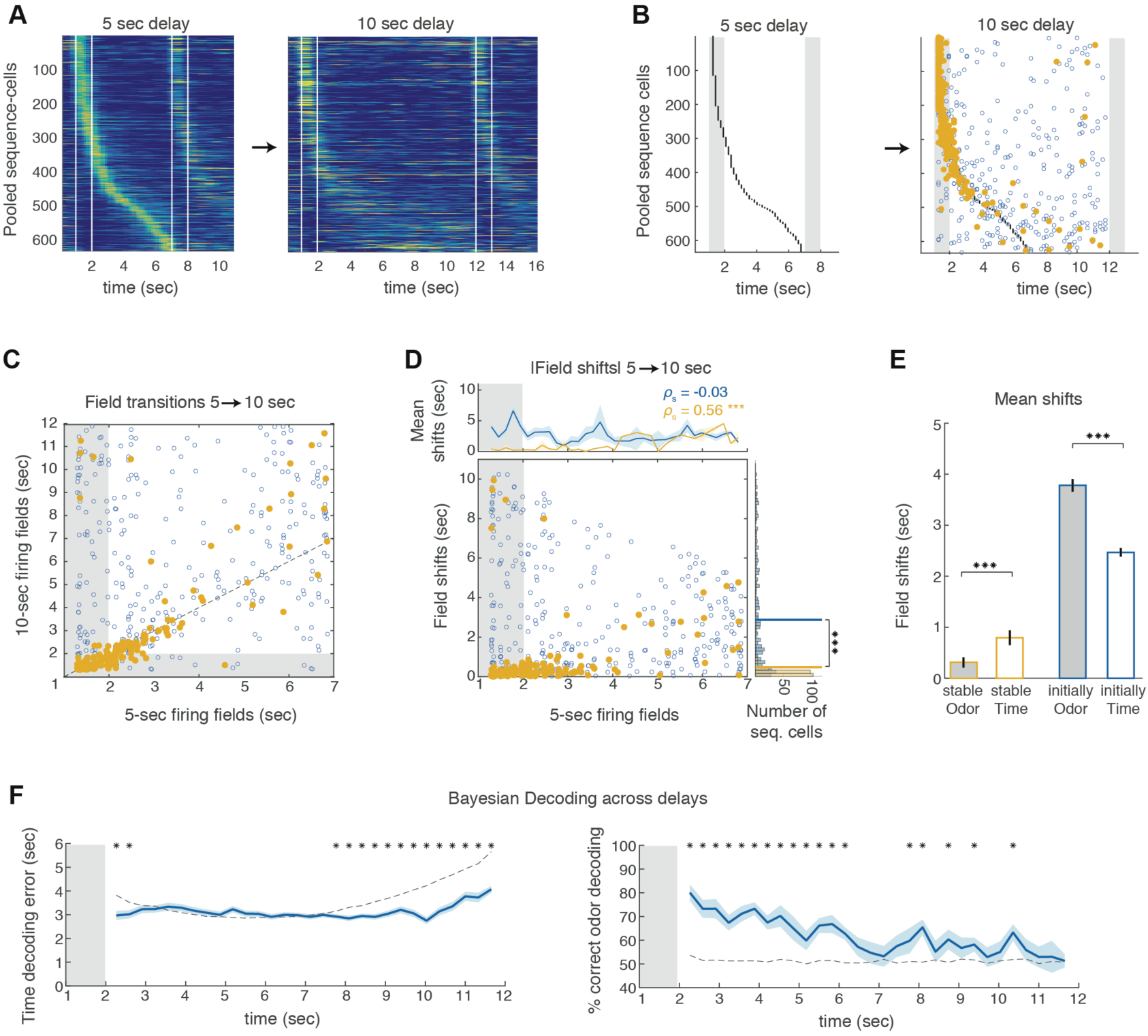
When extending the delay period, odor-cells retain their field whereas time-cells remap. **A.** Pooled sequence-cells from all recording days with 5 and 10 sec delays (N = 10 days). The 5 sec delay sessions were used to detect and stack sequence-cells (left). They are stacked in the same order over the 10 sec sessions (right). **B.** Same as **A** but showing only the locations of the sequence-cell fields (black dots) and the time-bin of their maximum mean firing rate at 10 sec delays (blue dots). Yellow dots: Significant fields at 10 sec sessions. **C.** Location of each cell’s maximum mean firing rate in 10 sec delay sessions as a function of their initial 5 sec delay field (blue circles). Yellow dots: Cells with siginifcant fields in 10 sec delay sessions. Dashed line indicates no change in firing field. Gaussian noise applied for plotting clarity of overlapping points, as in Figure 4F. **D.** Absolute time-shifts of sequence-cells between 5 and 10 sec delay sessions as a function of their 5 sec delay field. Top: Mean ± SE shifts as a function of a cell’s initial field. Colors as before. *ρ*s: Spearman correlation; asterisks: P < 0.001, permutation distribution. Right: Histogram of sequence-cells’ field shifts. Lines indicate distribution means. Asterisks: P < 0.001, two-sample Kolmogorov-Smirnov test. **E.** Mean ± SE shifts of odor-cells (grey bars) versus time-cells (white bars) for all common sequence-cells between 5 sec and 10 sec delay sessions (yellow; P < 0.001, WT) and sequence-cells without significant field in 10 sec sessions (blue outline; P < 0.001, two-sample t-test, Bonferroni over the two tests). **F.** Mean ± SE time prediction error (left) and trial-type prediction accuracy (right) as a function of time-bin, with Bayesian decoding of 10 sec delays, using the 5 sec delay rates to train (Methods). Asterisks and dashed curves: Significance and baseline as before (P < 0.05; WT; FDR).

Time-cells appeared to shift their field mostly forward in time, with stable ones forming an extended sequence when pooled over all sessions (Figure 5B-C). To test if this forward shift could suffice to decode timing information through their ensemble activity in extended delays, we trained a Bayesian decoder on the time-cell activity over the 5 sec trials and decoded time the 10 sec trials based on the same cells. Every other time-bin was decoded (so that firing rates for training and decoding had the same time span). This inherently assumes that sequences extend forward in a uniform way, so that e.g. the time-bin 8 sec in the extended delay can be decoded by training the decoder on the 4 sec activity of the same cells in the original delay. Following this scheme, we found that only time during the first ~1 sec and the final ~4 sec of extended delay could be efficiently decoded (Figure 5F; P < 0.05 versus chance; FDR), suggesting that, apart from the early time-cells that remained stable, late time-cells tended to shift forward in time so that they encoded the added period.

Finally, the delay in 10 sec trials was also tiled by temporal sequences. We asked what the activity of these new time-cells was in the 5 sec trials. By repeating our field-shift analysis in reverse, we noted that sequence-cells covering the additional delay time (last 5 sec) stemmed mainly from cells that tended to spike at the late stages of the original delay, though most of these cells did not exhibit a significant field there (Figure S6).

Collectively, these findings reveal that odor-cells retain their field during short-timescale adaptations, whereas time-cells readily remap or lose their field when the delay is extended, similarly to spatial representations in expanded environments^40^. Later time-cells were more prone to larger shifts and tended to cover the added delay. This dynamic adaptability of the temporal code compared to odor-cell stability does not support a simple feedforward mechanism, triggered by the odor-cells, and suggests that distinct mechanisms may drive the two representations.

### The number of time-cells increases during learning or the memory task, but not during passive exposure

Finally, we asked if the difference in stability of olfactory versus temporal representations is also reflected in how these two population dynamics are shaped? Is there a link between their emergence of either one and working memory activation during DNMS delays? Or, alternatively, do these two neural codes pre-exist even in the absence of a behavioral context?

To address these questions, we performed calcium imaging in mice (N = 8) while they were being trained to the full DNMS task. Most mice (N = 6) were imaged for 2 days during the pre-training stage (only non-match trials) followed by a series of consecutive imaging days in the training-stage (full DNMS), whereas the rest (N = 2) were imaged after 5 days in the training-stage. Typically, mice performed well in the pre-training stage (>80% hit rate on average), particularly on the second day of imaging. When first introduced to non-rewarded match trials in the training-stage, mice quickly learned to reject them, while retaining a high hit rate in non-match trials. Consequently, performance improved rapidly, mirroring the rejection rate increase (Figure 6A) and indicating a gradual increase in working-memory engagement. Most mice reached well-trained performance levels within 2-5 days of training and sustained them for the remainder of imaging days (Figure 6A).

**Figure 6:**
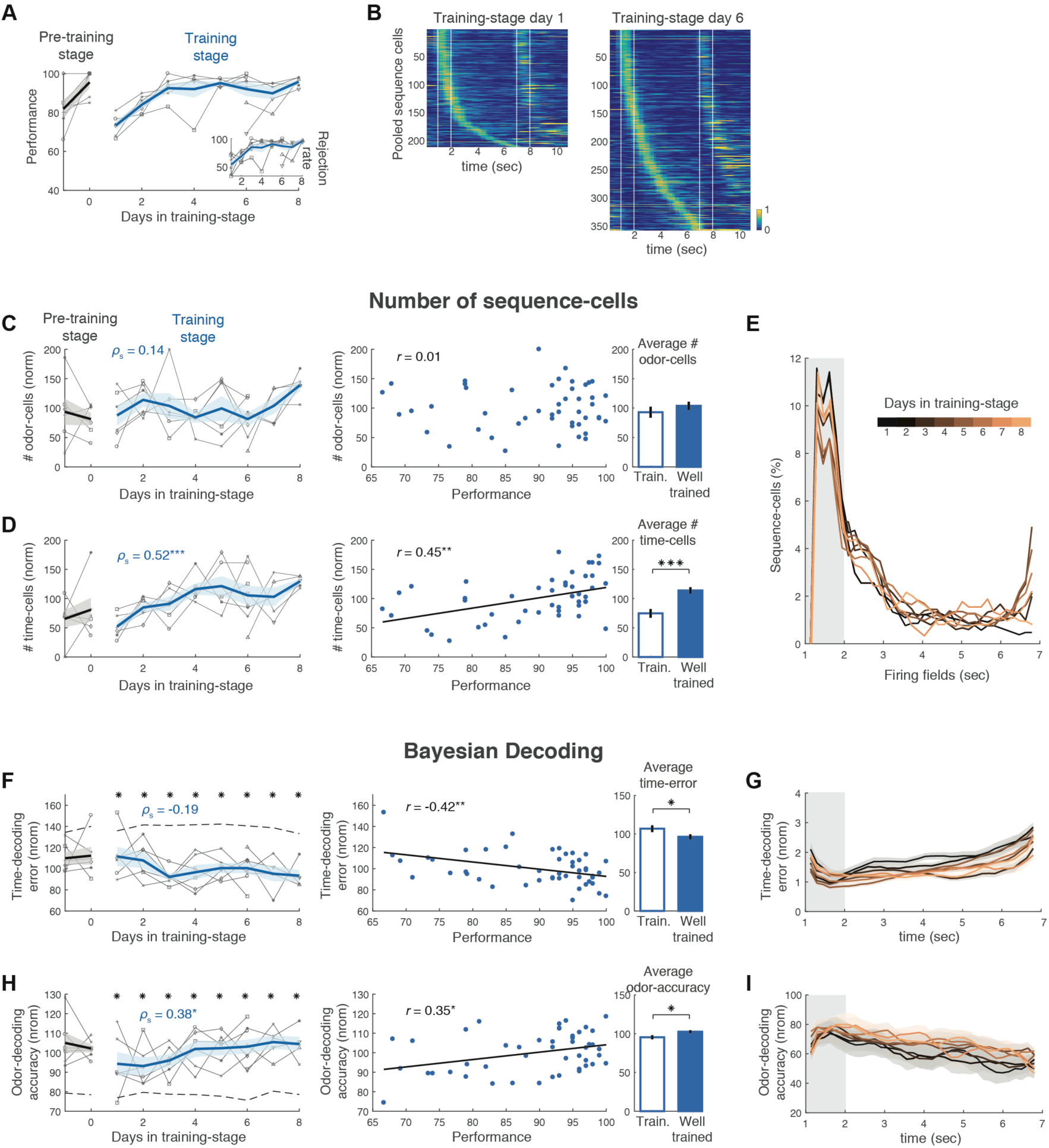
During learning of the DNMS task the number of odor-cells is stable while time-cells increase. **A.** Performance of individual mice (grey; N = 8) and mean ± SE (blue) as a function of consecutive days in the DNMS final training-stage. Two imaging days in the pre-training stage (only non-match trials) are also shown (black, N = 6). Inset: Rejection rate for match trials over days in the training-stage. **B.** Mean firing rates of all pooled sequence-cells in their preferred trials (N = 6 mice), recorded on the first day mice were on training-stage (left) and on the sixth day (right). The number of odor-cells is retained approximately fixed, whereas the number of time-cells increased dramatically. **C.** Left: Number of odor-cells per day, normalized over their mean number across all training-stage days for each mouse (shown in percentage), plotted as in **A**. *ρ_S_*: Spearman correlation. Middle: Normalized number of odor-cells versus each mouse’s daily mean performance. *r*: Pearson correlation. Right: Mean ± SE normalized number of odor-cells during all days at ‘training level’ (<90% mean performance) versus ‘well-trained’ level (≥90%). No significant correlations or differences are observed. **D.** Same as in **C** for number of time-cells, normalized as before. Time-cells increased monotonically over days (P < 0.001, Spearman correlation) and were significantly correlated with performance (P < 0.01, Pearson correlation; FDR over the two cell types; Black line: least-squares linear fit) so that their average number was significantly lower during training than in well-trained stage (P < 0.001; tailed two-sample t-test; FDR over the two cell types). **E.** Normalized distribution of fields per day in training-stage (smoothed with 5-point moving average). New time-cell fields are distributed similarly along the time-axis every day. **F.** Same as **C,D** for normalized Bayesian time-decoding error. **G.** Distribution of decoding-error over time-bins in each day. **H-I.** Same as **F-G** for normalized Bayesian odor-decoding-accuracy. Decoding improvement observed mostly at delay time-bins. Asterisks in left panels: Significant difference from shuffled baseline (dashed line; P < 0.001; WT; FDR).

We first examined whether learning of the task is linked with a change in the size of the cell assemblies generating either representation. Comparing pooled sequence-cells from the first day on training-stage versus the 6^th^ day (N = 6 mice) revealed that the number of odor-cells remained relatively stable while time-cells increased dramatically in number between these days (Figure 6B). Indeed, the number of odor-cells did not exhibit any trend over training days and no correlation with performance (Figure 6C; P > 0.05, Spearman and Pearson correlation respectively; permutation distribution test). In contrast, the number of time-cells per mouse was significantly correlated with training days (Figure 6D; P < 0.001, Spearman correlation; FDR over the two cell-types) as well as with mean performance per day (P < 0.01, Pearson correlation; FDR). Consequently, significantly less time-cells were found, on average, during days where mice were still learning the task (<90% mean performance), compared to when they were well-trained (>90%; Figure 6E; P < 0.001, WT; FDR over the two cell types). Interestingly, the distribution of time-fields did not change over days, retaining a similar power-law profile (Figure 6E), indicating that time-cells with early time-fields multiplied faster than those with later-fields.

Furthermore, as a result in the increase in the size of each temporal sequence, Bayesian decoding of both odor and time improved across days. Decoded-time error decreased during training, significantly anti-correlated with performance (Figure 6F), whereas decoded-odor accuracy significantly increased across days (P < 0.05, Spearman) in correlation with performance (Figure 6I; P < 0.05; Pearson correlation), leading to significantly lower mean time error and higher mean odor accuracy in well-trained days than training days (P < 0.05; WT; FDR). Due to the stable distribution of time-cells across days, this decoding improvement could be observed along delay time-bins each day (Figures 6G,I), while decoding efficiency over odor-bins was less affected.

To ensure that increased time-cells were not due to any general increase in dCA1 activity (e.g. due to increase in GCaMP6f expression), we applied the same analysis on the total number of ROI per day and the mean total firing rate and inter-spike intervals of cells during each trial. None of these measures exhibited any trend across days or any correlation with performance (Figure S7), suggesting that spiking activity across our field of view remained unchanged. Moreover, we repeated the analysis for the number of trials per day, excluding that more trials lead to higher time-cell detection. Finally, we also excluded motion on the treadmill as a confound, following the same procedure (Figure S7; P > 0.05 for all Spearman and Pearson correlations and WTs between training and well-trained numbers, in all tests on Figure S7).

We next asked whether task learning is linked with a change in the spiking properties of sequence-cells. Following the same analysis as before, we found no significant correlations of the selectivity index, activation probability or spiking variance of sequence cells with training days or performance (Figure S8; P > 0.05, same tests as above; FDR). Similar results were observed when examining odor- and time-cells separately (not shown). Conclusively, the spiking properties of sequence-cells remained stable, unrelated to performance.

Finally, we sought to ensure that the correlation between time-cells and performance was indeed driven by memory-mechanisms relating to learning the DNMS task, and not due to any increase in sequential spiking over days independently of working-memory context. We imaged naïve mice (N = 6) that were water deprived and habituated similarly to trained mice, but were completely naive to the DNMS task. These were imaged for 3 or 6 consecutive days, while passively exposed to a similar number of DNMS trials per day as trained. Sequence-cells were detected in these mice as well, with a similar distribution of fields and overall comparable spiking properties as in trained animals (albeit with a lower variance in spiking time, Figure S9). Interestingly, even though the percentage of ROI that was classified as odor-cells was similar to that in trained mice, these animals exhibited a significantly lower percentage of time-cells (Figure 7A; P < 0.01, WT, FDR). Moreover, unlike trained animals, pooled sequences on day 1 were similar to those on day 6 of DNMS exposure (Figure 7B). Indeed, the number of both odor- and time-cells did not change along imaging days (Figure 7C; P > 0.05), resulting in stable Bayesian decoding accuracy over days (Figure 7D; P > 0.05, Spearman correlation).

**Figure 7:**
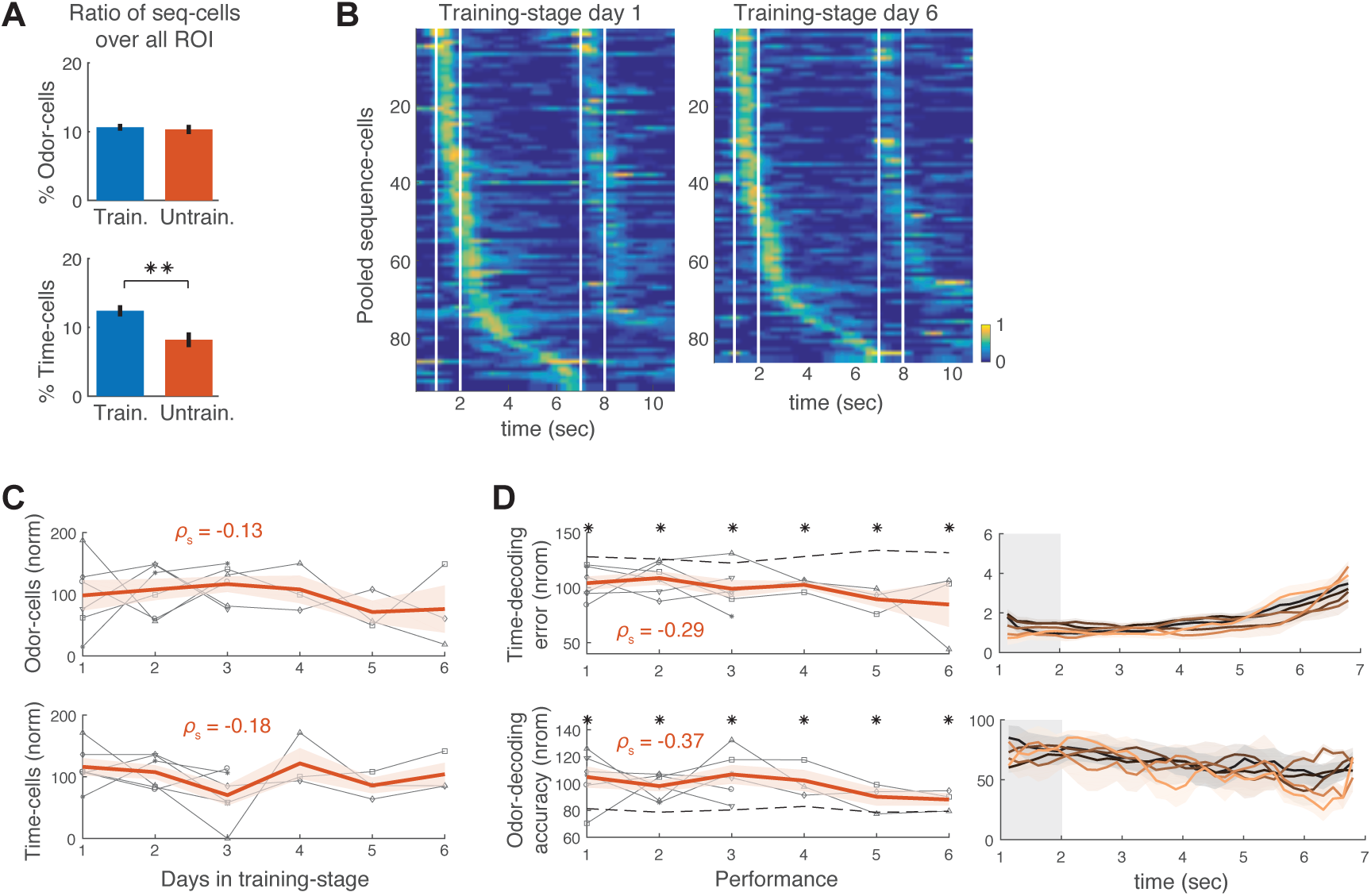
During passive exposure to DNMS, the number of time-cells does not change over days. **A.** Mean ± SE percentage of odor-cells (top) and time-cells (bottom) over all active ROI per day, in trained versus untrained animals (odor-cells: P > 0.05; time-cells: P < 0.01, right-tailed WT; FDR). Untrained animals (N = 6) were exposed to the full DNMS task (training-stage), similarly to trained ones. **B.** Mean firing rates of all pooled sequence-cells in their preferred trials, in untrained animals, recorded on the first day mice were exposed to the DNMS task (left) and on the sixth day (right). No particular change in numbers of sequence-cells was observed. **C.**, Normalized number of odor-cells (top) and time-cells (bottom) for untrained animals, plotted as in Figures 6**C,D**. **D.** Left: Normalized Bayesian time-decoding error (top) and odor-decoding accuracy (bottom) for untrained animals, plotted as in Figures 6**F,H**. Right: Distribution of decoding error and accuracy over time-bins as in Figures 6**G,I** respectively. No significant changes are observed in Bayesian decoding efficiency over days.

Our results suggest a link between temporal representations and learning of a memory-dependent task, while olfactory representations appear to be pre-existing in the CA1 network. Combined with our observed instability of time-cells across days (Figure 4), these findings indicate that a new set of time-cells is assigned to each sequence every day, with fields distributed similarly across the delay, yielding increased information encoding by the sequences. Interestingly, this phenomenon is linked with learning processes as it is not observed during passive exposure to DNMS trials.

## DISCUSSION

A recent model posits that hippocampal networks generate cell assemblies, spiking in sequence, as a means to organize and access neuronal representations of sensory experiences^5^. These sequences typically encode space, time, sensory or contextual information and may form a template for linking related memories^4^. However, each of these representations has been typically studied in isolation and across short time-scales. So, whether multi-modal representations have common or distinct emergence and stability characteristics remains unclear. Here, we employed two-photon calcium imaging *in vivo* on head-fixed mice while learning and performing an olfactory DNMS task, recording from the same dCA1 pyramidal populations over multiple days and delay durations. We observed (i) odor-cells, responding to olfactory stimuli that were followed by odor-specific time-cells, collectively forming spiking sequences that tile the entire delay period and are specific to the olfactory stimulus. (ii) Cells comprising either sequence were anatomically intermixed in the pyramidal layer. (iii) Odor was robustly represented by strongly activated cells, whereas time-information was sparsely encoded and progressively diminished throughout the delay period. (iv) Across multiple days, odor-cells retained stable firing fields whereas time-cells shifted their fields or lost them completely. (v) Similarly, when the delay period was extended within a session, odor-cells remained stable whereas time-fields remapped across the delay. (vi) During learning of the task, the number of odor-cells did not change, whereas time-cells increased as performance improved. This increase was linked to active learning as it was not observed during passive task exposure in untrained animals. Collectively, our findings reveal that memory-independent, stable and reliable olfactory representations, can coexist with memory-driven, sparse and highly dynamic stimulus-specific time-representations, spatially intermingled within CA1.

### A model of distinct mechanisms for stable olfactory and dynamic temporal representations in CA1

Comparative analyses of multi-modal representations can yield significant insights on the anatomical and neurophysiological mechanisms underlying their generation and evolution and their link to memory. Based on our collective findings, we propose that olfactory and temporal neural representations emerge through separate synaptic pathways and distinct neurophysiological mechanisms.

The striking stability of olfactory responses over days, modalities and learning, and their presence even in untrained animals imply that these are generated by preconfigured, strong, afferent synaptic pathways. Anatomically, only two synapses separate the olfactory bulb (OB) from CA1, since OB cells project to the lateral entorhinal cortex (LEC) which sends direct temporoammonic projections from LEC layer-3 to CA1 pyramidal cells^24^. These projections are necessary for olfactory associative learning^25^ and may underlie an increase in LEC-dCA1 coherence during olfactory-guided behavior^23^ and the increased excitability of CA1 cells during olfactory-discrimination learning^28^. OB output also reaches LEC via the piriform cortex which is involved in olfactory coding and rule learning^41,42^. These observations suggest that odor-identity representations may be generated outside the hippocampus, reaching CA1 pyramidal cells through LEC direct projections, forming stable odor-cells that are independent of memory-processing.

In contrast, the increase of time-cells during learning and the volatility of time-fields across days and within a session, when memory load is increased, suggests that temporal sequences are not preconfigured in CA1 networks but are generated by a memory-driven process. One possibility is that this process occurs internally, potentially within recurrent CA3 networks. Computational work suggests that such networks can self-organize and sustain multiple stimulus-specific assemblies, spiking in feedforward trajectories, through plasticity mechanisms^43^. CA3 neuronal populations do form behaviorally relevant spiking sequences^32,44^ and temporal-coding^13^ which could generate CA1 temporal codes through Schaffer collateral excitation. However, CA3 sustains more stable spatial mappings than CA1^37,45^ and this stability may extend to temporal sequences. In that case, CA3 assemblies would not adapt to increased delays fast enough to support the observed short-timescale instability in CA1. This remains to be explored by future research. Alternatively, a series of observations suggests that timing information in CA1 may be affected by activity in the medial entorhinal cortex (MEC). MEC neural populations form sequences while running in place on a treadmill^9^ and during timed immobility in a virtual navigation task^46^, signaling elapsed time. Furthermore, inactivating MEC output disrupts CA1 temporal coding and working memory performance, but not spatial and object coding^47^ or olfactory learning^25^. The same output controls temporal features in CA1 theta phase precession^48^ and replay activity^49^, further supporting a role for MEC projections shaping temporal codes in CA1. Yet, it still remains unclear whether MEC could produce context-specific timing or whether an interaction between the LEC and MEC activities or internal CA1 input summation may underlie the specificity of the CA1 temporal codes to the presented stimulus. Finally, entorhinal projections from either area are expected to function in conjunction with CA3 inputs and may be mediated by long-range inhibition^50,51^.

In summary, odor-cells may be driven by direct LEC temporoammonic projections to CA1, whereas time encoding may be shaped by MEC sequential activity. This scheme extends the established "what" and "where" information streams into the hippocampus^3,52^ to a stable "what" combined with a dynamic "what and when" information circuit, linked to learning and memory processing. It remains to be explored how these two information streams can achieve an impressive target-selectivity, such that one cell displays stable encoding for days whereas a neighboring cell can be highly dynamic even within a session.

### Expansion of temporal sequences

Input-specific spiking sequences form trajectories in a neural network’s state space, which can be robust to noise perturbations or small changes in initial conditions^31^, reliably conveying input information to downstream networks. The increase of time-cells during training supports this concept since it leads to improved cue and timing information but it is unclear which neural processes underlie it. Neuromodulatory inputs from the entorhinal cortex or septal regions may gate such dynamic changes. In the hippocampus, septal cholinergic inputs combined with CA3 excitation can drive long-term potentiation in CA1^53^, providing a plausible synaptic mechanism for the formation of spiking assemblies during the delay, as seen in visual cortex^54^.

The fact that some time-representation existed even in untrained animals supports recent observations of hippocampal sequences arising even in the absence of external cues^55^. However, since odor stimuli were not associated with reward, no relevant information needed to be conveyed during the delay period. Therefore, the mechanisms responsible for increasing the temporal representations were not at play in this case, leading to low information-content sequences, consisting of few cells that remained unchanged.

### Stability of representations

The study of the stability of neuronal dynamics over long timescales has only recently become possible through the advancement of optical imaging methods. Both stable and dynamic representations have been reported in cortical areas^56,57^. Research on hippocampal activity has also pointed to both stable encoding^30^ as well as dynamic spatial^37,38^ and fear representations^39^ drifting to different neuronal assemblies over days. The gradual drift in temporal coding in our data complements that latter framework. Interestingly, despite this gradual drift, temporal and stimulus information can still be extracted from the same population after multiple days, which corroborates recent findings of non-memory-related CA1 time-encoding drifting across three days^58^.

Conversely, olfactory representations exhibited striking stability during learning and over multiple days, leading to robustly encoded odor-information that can be relayed downstream. Similar stability has been reported for sensory representations in cortical networks^56^ or for spatial representations in the dentate gyrus^45^, but not in CA1. It remains to be explored whether it is specific to olfactory inputs or extends to other sensory modalities^18,20^, and whether it is specific to working-memory cues or common across behavioral paradigms.

### Links to working memory

Working memory is thought to incorporate a large network of cortical and subcortical structures, including the parietal cortex^14^, entorhinal cortex^59^ and hippocampus^60^, with prefrontal-hippocampal synchrony being a key component for active memory maintenance^60^. Our observed time-sequences are odor-specific, in agreement with previous studies showing distinct temporal codes for different memories^11,12^. Moreover, the time-cell increase during learning supports previous observations that temporal coding develops in conjunction with temporally structured memories^10,11^. These characteristics suggest a link between working memory and hippocampal time codes. Information carried by time-cells, may be conveyed to outside the hippocampus, through e.g. polysynaptic and direct projections to the OB^62^ and olfactory cortex^26^ or monosynaptic projections to the medial prefrontal cortex which is involved in learning the DNMS task^34^ and exhibits similar temporal codes^15^. Sequence expansion during learning implies increased information sent out to such areas, which may be important for active memory retention or rule learning during the training process.

### Future directions

Untangling the cellular and network mechanisms that generate these combined representations, how they are linked to learning and memory and what information they relay to downstream regions is crucial for understanding the role of hippocampal population dynamics in memory formation. Future studies will be needed to dissect the role of projections from the LEC, MEC, and CA3 for the emergence, stability and plasticity of time and sensory representations. Furthermore, it will be critical to study the role of specific interneuron types in sculpting sensory and time representations and regulating during learning. The basic rules governing the emergence and plasticity of internally generated sequential activity in hippocampus will likely apply to multiple other structures engaged by the maintenance of information in the brain.

## ACKNOWLEDGEMENTS

We dedicate this work to the late Dr. Howard Eichenbaum whose research inspired it. We are very grateful to J. Sadik, N. Saboori, M. Bedrossian, B. Nosrati, T. Taimoorazy and Y.M. Chan for their valuable help in training animals and for feedback on training protocol. We thank K. Maguire, T. Shuman, D. Aharoni, S.W. Hur and A. Bellafard for technical support. We also thank D. Buonomano and W. Hong for valuable feedback on this manuscript. This work was supported by the US National Institutes of Health (R01 MH101198, RO1 NS099137, RO1 NS090930, R01 MH105427), the Intellectual and Developmental Disabilities Research Center at UCLA (U54 HD87101) and a Veterans Affairs Merit Review Award (1I01BX001524-01A1).

## AUTHOR CONTRIBUTIONS

J.T. and P.G. conceived and designed the experiments. J.T. constructed training and experimental rigs, created software for behavioral training, conducted experiments and analyzed experimental data. E.P. and J.T. created software to analyze calcium imaging data. J.T. and A.M. acquired experimental data. J.T., A.M., J.A., K.S., E.H. prepared experiments. J.T. and P.G. wrote the manuscript.

## COMPETING INTERESTS STATEMENT

The authors declare no competing financial interests.

## DATA AVAILABILITY STATEMENT

All datasets generated during the current study and all computational codes developed for analyses are available from the corresponding authors on reasonable request.

## METHODS

### Animals

A total of 14 adult male mice (11-31 weeks old) were used for *in vivo* two-photon calcium imaging experiments: 7 Gad2-Cre:Ai9 mice (Gad2^tm2(cre)Zjh^/J crossed with B6.Cg-Gt(ROSA)26Sor^tm9(CAG-tdTomato)Hze^/J), 5 Gad2-Cre:Ai14 (Gad2^tm2(cre)Zjh^/J crossed with B6.Cg-Gt(ROSA)26Sor^tm14(CAG-tdTomato)Hze^/J) and 2 Gad2-Cre mice. An additional 10 adult male mice (13-25 weeks) were used for behavioral control experiments: 2 Gad2-Cre:Ai32 (Gad2^tm2(cre)Zjh^/J crossed with B6;129S-Gt(ROSA)26Sor^tm32(CAG-COP4*H134R/EYFP)Hze^/J), 2 Gad2-Cre:Ai9 and 6 C57B/6J. All animals were acquired from The Jackson Laboratory and were group housed (2-5 per cage) on a 12 h light/dark cycle. All experimental protocols were approved by the Chancellor’s Animal Research Committee of the University of California, Los Angeles, in accordance with NIH guidelines.

### Surgical Procedures

Mice were anaesthetized with isoflurane (3–5% induction, 1.5% maintenance), their scalp was shaved and they were placed into a stereotactic frame (David Kopf Instruments, Tujunga, CA) on a feedback-controlled heating pad (Harvard Apparatus) to maintain body temperature at 37°C. Eyes were protected from desiccation using artificial tear ointment. The scalp was sterilized with betadine and infiltrated with lidocaine (2%; Akorn, Lake Forest, Illinois) as a local anesthetic. An incision was made to expose the skull from bregma to lambda. Fascia was removed by applying hydrogen peroxide and the skull was stereotactically aligned. A small burr hole was made on the right hemisphere (−2mm posterior and 1.8 mm lateral to bregma), using a dental drill. Using a Nanoject II microinjector (Drummond Scientific), 1500 nl of 1:10 saline-diluted AAV1.Syn.GCaMP6f.WPRE.SV40 virus (diluted immediately prior to surgery; titre: 4.65 × 1013 GC/mL; Penn Vector Core) were injected into dorsal CA1 (1.3 mm ventral from dura) at a 50-100 nl per minute. 60 minutes after the termination of viral injection, a circular craniotomy (3mm diameter) was made around the injection site. Dura over the exposed brain surface was removed and the cortical tissue above the dorsal CA1 was carefully aspirated using a 27 gauge blunt needle. Buffered artificial cerebrospinal fluid (7.888g NaCl, 0.372g KCl, 1.192g HEPES, 0.264g CaCl2, 0.204g MgCl2 per 1000 ml milipore water) was constantly applied throughout the aspiration to prevent desiccation of the tissue. The aspiration ceased after partial removal of the corpus callosum and bleeding terminated, at which point a 3-mm titanium ring with a glass coverslip attached to its bottom was implanted into the aspirated area and its circular flange was secured to the skull surface using vetbond (3M). A custom-made lightweight metal head holder (headbar) was attached to the skull posterior to the implant. Cyanoacrylate glue and black dental cement (Ortho-Jet, Lang Dental) were used to seal and cover the exposed skull. Mice that were used for behavioral experiments were only implanted with a headbar and were not given a craniotomy or aspiration. During recovery (~7 days) mice were administered carprofen (5 mg per kg of body weight) for 3 days as a systemic analgesic and amoxicillin antibiotic (0.25 mg ml^-1^ in drinking water) through the water supply for 5 days. Their weight was monitored daily.

### Experimental Setup

The treadmill consisted of an 8-inch Styrofoam ball (Graham Sweet) suspended through a metal axis, allowing for 1D rotation. It was housed within a dark enclosure. Mice were headfixed on the treadmill by attaching the implanted headbar to a custom-made metal holder. Locomotion was recorded as an independent analog signal, using a custom printed circuit board based on a high sensitivity gaming mouse sensor (Avago ADNS-9500), connected to a microcontroller (Atmel Atmega328) sending data to a dual channel digital-to-analog converter at 100 Sps.

A constant stream of clear air (~1 L/min) was transferred to the behavioral rig via Tygon PVC clear tubing and was supplied to the mouse through a custom-made lickport. During odor stimulation a dual synchronous 3-way valve (NResearch), placed ~6 cm away from the mouse, would switch from the clear air stream to the odorized one for 1 sec. Odorized air was created using a 4-ports olfactometer (Rev. 7c; Biology Electronics, Caltech), supplying air to either of two glass vials containing liquid isoamyl acetate (70% isoamyl acetate basis, FCC; Sigma Aldrich) or pinene ((−)-α-Pinene, ≥97%, FCC; Sigma Aldrich) odorants, diluted in mineral oil at 5% concentration (unless stated otherwise). For the non-match-to-long-duration-sample task (NMLS) experiments, all 4 ports were connected to odor vials (see below). Odorized air from each vial would reach the behavioral rig through separate Tygon tubing each leading to a 3-way solenoid valve (Lee Company). For olfactory stimulation, the corresponding olfactometer port started supplying air to its attached odor vial 1 sec prior to actual stimulation to allow for odorized air to travel through the tubing and reach its corresponding solenoid valve. During the 1 sec stimulation, that solenoid turned on allowing the corresponding odorized air to enter a short common path for both odorants (~4cm) ending at the dual synchronous 3-way valve and from there being released to the mouse through the lickport (~10 cm total common path for the two types of odorized air). At the offset of the stimulus, the solenoid turned off and the dual 3-way valve switched the airstream back to clear air, ensuring a constant flow of air to the mouse and a quick clearing of the odorant from the air around the mouse. The odorized and clean air were set to similar airflow values (~1 lt/min), measured with a flowmeter (AWM3300; Honeywell).

Odorant concentration in open air was measured through a mini photoionization detector (PID; 200B mini PID; Aurora Scientific), located at the distance of the mouse snout from the air-delivering lickport. Lower ionization threshold for pinene (odor-B) lead to higher deflections in the PID signal during its presentation compared to isoamyl acetate (odor-A). Therefore, potential odor lingering in the small common tube path, during the delay, and mixing with the second odor would result in a larger PID deflection during the second odor in B-A trials (lingering pinene molecules mixing with isoamyl acetate) than in A-A trials (lingering isoamyl acetate mixing with itself). This was observed when creating a very long common path length, but not in our default set up (Figure 1S, indicating that no odor mixing is created during the second odor.

Licking was detected using a battery operated, custom-made, printed circuit board operating as a lickometer^63^. One end of the circuit was attached to the metal headbar on the mouse and the other to the metal tube delivering water. Whenever the mouse would lick the tube, an electrical circuit would close, creating a voltage drop that was recorded as a continuous RSE analog signal. Water droplets (~10 µl) were released by a 3-way solenoid valve (Lee Company) and delivered to the mouse through a metal tube on the lickport. At the end of each trial, after the response window, vacuum was applied for 3 sec, through a metal tube underneath the water delivering tube, to clear any lingering water and assist in removing any odorized air.

The behavioral rig was controlled with custom written software (Matlab) and through a data acquisition board (USB-6341; National Instruments).

### Behavioral training and recording protocol

After the recovery period, mice were briefly handled for two days and water restriction was initiated. They were provided with 1-1.5ml of water daily, either throughout or post training. Their weight was maintained at around 85% of their final weight pre-water restriction initiation. Mice were initially habituated to head fixation on the spherical treadmill for 3 consecutive days. On the second day, the lickport was placed in its normal position for habituation to the metal ports and air flow. Mice were given water drops through the lickport on the third day of habituation.

DNMS training consisted of 3-4 days of ‘shaping-stage^’43^ where mice were presented with 20-trial sessions of non-match trials and the water reward was delivered automatically at the start of the response window in each trial. This was followed by a ‘pre-training’ stage for 2-4 days where mice were presented with non-match trials but had to actively lick the lickport during the response window to release the water drop. They were moved to the final training stage once they performed at >85% hit trials per day. In the ‘training stage’, mice were presented with the full task in a series of sessions per day. Each session again consisted of 20 trials and match trials were intermingled with non-match ones with each group of 4 consecutive trials containing all possible 4 combinations of odors in random order. Mice were not punished if they licked at match trials but no reward was presented. Within ~2-5 days they learned to refrain from licking during match-trials. During training, mice gradually refrained from licking during the delay period as well, and well-trained mice would typically initiate licking during or right after the second odor. Responses were assessed based on licking during the response window only. If any licking occurred during that window, the trial was a hit (non-match trials) or false alarm (match trials) accordingly. In the opposite case, they were labeled as miss or correct rejection respectively. Performance over each behavioral session, was quantified as the percentage of hits and correct rejections over the 20 trials. Daily performance was the corresponding ratio over the total number of trials (all sessions) during that day. Mice were typically presented with 5-7 sessions daily.

Two-photon imaging sessions for mice under training started as soon as they were ready to move from the ‘pre-training’ stage to ‘training stage’ (N = 6 mice), at ~2-3 weeks post-surgery. They were first imaged for 2 days while performing ‘pre-training’ behavioral sessions (non-match trials only). 3 of the mice received 3 ‘pre-training’ sessions followed by 3 training sessions on their second day of imaging. The other 3 received 5 pre-training sessions on their second imaging day and were moved to training sessions on their third day of imaging. Finally, 2 mice were imaged only after they had received 5 days of training stage (not having reached well-trained level yet) and they were only imaged at training stage.

Some mice (N = 6) were imaged while naïve to the task. Only the 3 days of habituation were applied (without any water droplets delivered on day 3 of habituation) before onset of calcium imaging, upon which they were presented with normal DNMS sessions for 3 or 6 consecutive days (N = 3 in each case), same as with trained mice. 5/6 naïve mice never licked the lickports for a reward. One naïve mouse kept licking the tubes and received a series of water rewards, but only early sessions (3 days) where the mouse had not grasped the task were included here. These animals were imaged 2-3 weeks post-surgery to match the time the trained animals were imaged and to allow for GCamp6f to be expressed.

### Behavioral control experiments

Non-match-to-long-duration-sample task training (NMLS; N = 3 mice) consisted of the same training steps as DNMS, but the first odor was presented throughout a 5-sec period and was immediately followed by the second odor. Varying odorant concentration vials were used only for the first odor whereas the second odor was delivered by different olfactometer ports connected with 5% odorant vials. For experiments with odors turned off, the olfactometer ports were deactivated, halting any flow of odorized air, but trials were otherwise normal. For experiments with variable airflow, the airflow for both odor-ports on the olfactometer was altered randomly for each port at the beginning of each behavioral session, between ~0.4 −1 lt/min, measured by the flowmeter.

### In vivo two-photon imaging

A resonant scanning two-photon microscope (Scientifica) was used for calcium imaging, recording 512×512 pixel frames at 30.9 Hz, with a 16x 0.80 NA objective (Nikon) yielding a 500×500 µm field of view. Excitation light was delivered with a Ti:sapphire excitation laser (Chameleon Ultra II, Coherent), operated at 920 nm. GCamp6f and Td-Tomato fluorescence were recorded with green and red channel gallium arsenide photomultiplier tubes respectively (GaAsP PMTs; Hamamatsu). Microscope control and image acquisition were performed using LabView-based software (SciScan). Imaging and behavioral data were synchronized by recording TTL pulses, generated at the onset of each imaging frame, as well as olfactory stimulation digital signals at 1 kHz, using the WinEDR software (Strathclyde Electrophysiology Software). A single fixed field of view was imaged every day for each mouse. Laser power was kept to a minimum and no photo-bleaching was observed.

### Calcium imaging data processing

Data was pre-processed in MATLAB using a custom-built pipeline based on the CaImAn package^36^ for large scale calcium imaging data analysis. To reliably track neurons over multiple days in the presence of changes in the field of view, active neurons were extracted separately for each day and were then registered against each other.

#### Motion correction and source extraction

Datasets from each day where corrected for motion artifacts using online piecewise rigid registration^64^, where the template obtained during motion correction of session X was used to align data from session X+1. Data from all the sessions each day was then concatenated and downsampled in time by a factor of 5 to increase the signal-to-noise ratio. The spatial footprints of the active sources (ROI) were then extracted from the downsampled data using an implementation of the CNMF algorithm in spatial patches^35,36^. Spatial and temporal correlation thresholds for ROI detection were set to 0.6 and 0.8 respectively, acceptable ROI size was limited to 50-150 pixels, and an ROI eccentricity threshold was set at 0.97 to avoid elongated ROI that were axodendritic segments. These values were determined after examination of the relationship between these thresholds and the final accepted ROI spatial footprints and temporal traces, using a custom graphical user interface on a set of our recordings (Figure 1D). The spatial footprints were then used to obtain the traces at the original frame rate by solving a non-negative regression problem. The computed traces were first transformed in ΔF/F units and were then deconvolved using the OASIS algorithm for fast nonnegative deconvolution^65^, separately for each session, to correct for possible bleaching artifacts or variation of laser power between sessions. Noise levels were computed for each ROI using a power spectral method, and spike thresholds were set to 3 x noise level, except for two animals with low signal-to-noise ratio and sparse activity, were the threshold was lowered to 1.5. Finally, the deconvolved traces were binned in time using 50% overlapping 320 ms time bins (Figure 1E). This signal is used as a proxy of spiking activity per unit of time and is referred to as ‘spiking rate’ throughout the text. This pipeline generated a set of spatial footprints and temporal traces (both in ΔF/F unit and in deconvolved neural activity) for each animal at each day of recordings.

#### Inhibitory neuron detection and removal

Inhibitory neurons were identified based on their static Td-Tomato fluorescence recorded for 500 frames at the beginning of each imaging day on the red-channel PMT, together with the functional (green) channel. The red channel was first aligned to the green one by using the motion displacement field that was estimated during the motion correction of the green channel. Then the average of the red channel was computed and the resulting image was segmented to obtain contour plots of the inhibitory neurons. The segmentation was performed by using adaptive thresholding (to model different illumination levels within the field of view), with a threshold at each location computed as the Gaussian weighted average of a small neighborhood around the location. The resulting thresholded image was segmented using connected components. For each connected component, the corresponding part in the mean image was then thresholded at 75% of its maximum value, and after this thresholding and another round of connected components analysis, components of small size (<30 pixels) or odd shape (>0.95 eccentricity) were removed. The remaining components identified the locations and spatial extent of the inhibitory neurons within the field of view (Figure 1D). These components were then registered with the CNMF identified components from the functional channel using the procedure described above. Components that were matched corresponded to inhibitory neurons that were active during this imaging session, whereas mismatched components corresponded to inhibitory neurons that were either silent or whose activity did not meet detection criteria by CNMF (slow traces potentially due to prolonged high-frequency spiking). Matched components were removed from the final pool of ROI, so that only pyramidal cell activity from the green channel was analyzed further.

#### Pairwise Registration

To register components across two different days we followed the following procedure: First the ROI from both days were aligned to the same FOV, by computing a motion transformation between the templates from day 1 to day 2 and using this transformation to align the components from day 1 to the FOV of day 2. Let 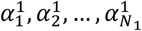 and 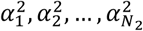 be the set of (aligned) spatial footprints from the two sessions. The components were first 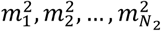 transformed into binary masks 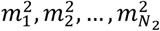 respectively, by thresholding each component at the 10% of its maximum value. An intersection over union metric was used to quantify the distances between the footprints from the different days:

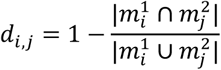

Based on this metric a matrix of pairwise distances was constructed. Distances between components where the one was a subset of the other (>60% pixels of smallest of the 2 ROIs overlapping with those of the largest) were set to zero, whereas high distances *d*_*i,j*_ > 0.98 were set to infinity. Components were then registered to each other using the Hungarian algorithm (a polynomial time algorithm for solving the linear assignment problem). Setting weakly overlapping components to have infinite distance prevented false pairings between components which were instead left unmatched between these two particular sessions.

### Calcium and Behavioral Data Analysis

All analysis was performed on the binned spiking rate traces from all pyramidal cells each recording day, using custom written analysis software (Matlab).

#### Sequence-cell detection and analysis

Odor and time-cells were detected separately for all A-trials and B-trials (initiated by odor-A or odor-B respectively) each day for each cell, through the following process: First we computed its average firing rate for the corresponding group of trials within the time interval from the first odor onset up to the second odor onset (‘odor-delay interval’). The maximum average rate was computed together and the time-bin it occurred in was considered the cell’s ‘firing field’. Only if the cell spiked within that interval at least in 10% or 3 (whichever was greatest) trials of the corresponding type was considered further. Its firing rate trace over the odor-delay interval was then circularly shifted by a random interval up to ± ½ x odor-delay interval, separately for each trial of that type and the maximum mean firing rate over the shifted trials was computed. This process was repeated 1000 times, generating a distribution of maximal rate values. A cell was considered to have a significant firing field if its maximum mean firing rate at that time-bin within the odor-delay interval was larger than the 95^th^ percentile of that shuffled distribution. If the time bin of that maximal rate was during the odor stimulation, it was considered an odor-cell, otherwise a time-cell.

We computed the selectivity index (*SI*) of each sequence cell as:

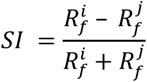

Where 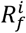 is the cell’s mean firing rate at its firing field *f* over all *i*-trials, whereas 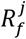 is its mean firing rate at the same time bin but over the opposite type of trials *j* (with *i* and *j* representing odor-A or odor-B accordingly). Cells with a negative SI over the trial type under consideration were discarded from the corresponding sequence (as preferring opposite trials).

For the three mice receiving a combination of 3 ‘pre-training’ and 3 ‘training-sessions’ on their second imaging day (see above) this process was performed separately for each training stage. For experiments where the odors were turned off after the initial sessions, the sequence cells were computed using only the trials with the odors on. For trials with variable odorant concentrations, they were computed based on all trials. Finally, for imaging days with multiple delays, sequence cells were detected separately over trials of each delay duration.

The ‘participation’ of each cell to its sequence was quantified as the percentage of corresponding trials where the cell had a non-zero firing rate inside its firing field. Spiking variance was quantified as the variance of each trial’s maximal firing rate time bin over all trials of a given type. These two measures, as well as the SI, are always plotted smoothed with a 5-point moving average.

For plotting, mean firing rates of each sequence cell were normalized by their maximal firing rate at their firing field. The same normalization was applied for both trial types.

For spatial distribution analysis, the centroid of each sequence-cell’s spatial footprint was computed. All pairwise centroid Euclidean distances were measured as well as pairwise time distances between firing fields for all time-cells.

In Figure 3A, the mean place-field distribution was approximated by a two-term power law distribution:

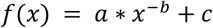

with a = 27.06, b = −2.68, and c = 0.92, using the trust region reflective algorithm (*fit* function in Matlab). The goodness of fit was assessed by creating random sampled distributions from this power law, of equal size to the original, with 2500 repetitions, fitting each sampled distribution with the same algorithm, and comparing the Kolmogorov-Smirnov (KS) statistic of the original distribution with those from the random sampling^66^. The P-value corresponds to the ratio of distributions with KS statistic lower than the default distribution.

#### Motion Analysis

The locomotion signal (see above) had its baseline removed (mode value), was binned in the same manner as for the deconvolved calcium traces and its summed value over each trial was computed. Trials were ranked based on the total motion measure and 30% of trials with lowest locomotion were compared with 30% of trials with highest locomotion (Figure S3).

#### Analysis Across Multiple Days

ROI were matched across all pairs of imaging days per mouse. ‘Recurring sequence-cells’ were defined as two matched ROI that had a significant field in the same sequence both days. ‘Stable cells’ were detected by using the last imaging day as reference and tracking which ROI of that day had a match for all other days. The same method was used to find ‘stable sequence-cells’, again using the final day’s sequence cells as reference and tracking those that had a match with a cell belonging to the same sequence each other day. To compute field shifts across two consecutive days (Day 1 and 2), we took all sequence cells from Day 1 that were matched to an ROI on Day 2 and computed the maximal firing rate time bin of that matched ROI across the corresponding trials of Day 2 (along the odor-delay interval as usual). Field shifts of recurring sequence-cells were those where the matched ROI was a sequence cell for that trial type as well. If the matching ROI never spiked on Day 2 in the corresponding trials, the cells were discarded from field-shift analysis. When plotting average sequence firing rates (color plots; Figure 4D), each cell’s rate was normalized by its mean rate in its field on Day 1.

#### Analysis Across Multiple Delays

A set of well-trained mice (N = 5) were imaged during both 5 and 10 sec sessions on the same day. Typically, these mice would receive a set of 2-3 behavioral sessions with 5 sec delay followed by the same number of sessions with 10 sec delays. Each animal was exposed to extended delays for 2-3 days in total. Two animals received a sequence of sessions that dropped back to 5 sec after the delay extension to 10 sec (during 1 day for one animal and 2 days for the other). Removing these imaging days where the delay was reduced again, did not affect our results.

Separate sequence-cells were detected each day over the 5 sec trials and the 10 sec trials. ‘Stable cells’ refer to those that have a significant firing field in the same sequence in both delays (not necessarily in the same time bin thought), whereas ‘unstable’ cells refer to cells that have a field in one delay but not the other. As in multi-day analysis, for unstable cells we use their maximal average firing rate location over their preferred trials as proxy of a field. If a sequence-cell never spiked in the other time-type trial it was discarded from field-shift analysis. When plotting average sequence firing rates (color plots; Figure 5A), each cell’s rate was normalized by its mean rate in its default delay.

#### Analysis over Training Days

Performance was calculated separately for each behavioral session (20 trials) and was averaged over all sessions per imaging day. Rejection rate was quantified as the ratio of match-trials where the mouse did not lick and is only defined for the training-stage trials (full DNMS). To pool individual mice, data from each mouse (number of cells, Bayesian decoding measures etc.) were normalized over their mean number across all training-stage days, and are plotted as percentages. The same normalization was applied to data points in pre-training stage.

Mean total rate was computed by summing the deconvolved calcium trace of each cell over all time-bins during each trial, and averaging over all trials and all cells. Mean motion was computed by summing the motion signal (see above) over all time bins of each trial and averaging over all trials. Inter-spike intervals (ISI) were computed at each trial by making the deconvolved calcium trace of each cell binary (turned to 1 for any value > 0) and computing the time interval between two consecutive 1s following 0s. Mean ISI was computed by averaging each cell’s ISI over each trial, and taking the mean over all trials and all cells.

#### Principal Components Analysis

Principal component analysis (PCA) was applied to the collected firing rates of sequence-cells in each imaging day. Each cell’s rate was z-scored over all trials of a given odor and the PCA scores were computed. The explained variances for each PC were computed and PCs explaining >80% mean cumulative variance were kept, separately for each trial type. If that threshold was never reached, then all PCs were kept. The minimum number of the two PC limits was finally applied to both trial types. Trajectories along the multidimensional space of those PCs were computed for each trial and the mean trajectories over the two trial types were kept. The Euclidean distance between those was computed and smoothed with a 0.65 sec-window moving average. Baseline distances were computed for each session by circularly shifting the rates of sequence-cells along time by a random interval up to ± ½ x trial duration with 100 repetitions and repeating the PCA analysis for each repetition.

Plotted PCA trajectories in Figure 3C were smoothed over the 3 PC space with spline interpolation and color coded using linear interpolation over time.

#### Bayesian Decoding

A Bayesian decoder was used to assess how well can collective sequence-cell activity predict time during the trial, as well as trial-type^67^. A separate decoded was constructed for each imaging day, using the firing rates of all sequence-cells (both sequences pooled), over the odor-delay time interval. Only correct trials out of the first 2/3 of all trials of the day were used for training the decoder and the last 1/3 trials were used for decoding. Both correct and error trials were decoded, unless otherwise stated.

Each cell’s mean firing rate over each trial-type was smoothed using 1sec-window moving average. In order to decode the trial-type (odor presented), we considered time space to be 2 x odor-delay interval (12 sec long in total). We thus concatenated along the time axis, each cell’s mean rate over the two trial-types. The decoder, trained by these concatenated firing rates (mean rate over A-trials followed by mean rate over B-trials) would thus predict a time point along the doubled time interval 0-12 sec. If the time point was within the first half (0-6 sec) it corresponded to that time point of an A-trial, whereas if it was within the second half (6-12 sec) it corresponded to the analogous time point of a B-trial.

Assuming Poisson distributed spiking and statistical independence of sequence cells, the decoded time point 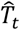 from activity at time bin *t* is given by:

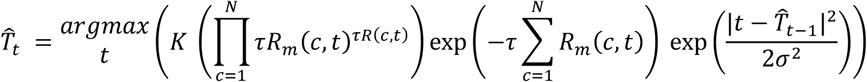

where *c* = 1,…,N are the pooled sequence-cells and *R_m_(c,t)* is the mean firing rate of cell *c* at time bin *t* (with bins concatenated over both trial types, spanning 12 sec) over all training trials. *R(c,t)* is the corresponding firing rate at the decoded trial, τ is the bin duration and *K* is the probability of being at time bin *t* of a particular trial type which is a constant, proportional to the ratio of trials of that type over all training trials^67^. The last term functions as a continuity constraint, limiting the decoded time bin to be in a relative proximity to the previous one^67^, with σ = 3 sec. Time bins with no activity from any sequence cell were not decoded. Chance baselines were computed by randomly shuffling the cell identities for each decoded trial with 1000 repetitions and decoding the shuffled cells for each repetition. Time prediction error refers to the mean absolute time distance between a given timepoint from either trial-types and the decoded one: 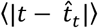, and does not take the trial-type into account. It thus functions as a measure of the time-information carried by the sequence-cells. Odor prediction accuracy refers to the percentage of correct trial-type decoding at any given timepoint (i.e. correct half of extended time-axis) and functions as a measure of the information on the odor identity carried by the sequence cells. Both measures were averaged over pooled trials from all analyzed sessions, for each decoded time point separately.

For cross-days analysis, we applied the Bayesian decoder on the firing rates of the recurring sequence-cells between each pair of days. Day pairs with no common sequence cells were not processed. All correct trials from Day 1 were used to train the decoder and all trials of Day 2 were decoded. Baseline was created separately for each day pair, by randomly shuffling the identities of the recurring cells in Day 2, with 1000 repetitions, and applying to decoder at each repetition. Decoded-time errors over each time point were computed for each day pair, concatenated for both trial types. Errors from all day pairs of any given distance were pooled together and their mean value for that given day distance was computed and plotted (Figure 4I). The same process was applied to odor-decoding accuracy as well as the corresponding baselines for both measures. Data points from zero distance between days in Figure 4I correspond to decoding all trials from a single day using the same trials (subset of correct trials) for training the decoder.

For multi-delay analysis, the Bayesian decoder was trained only with the firing rates of 5 sec-session time-cells and was used to decode odor/time through the activity of these cells during the 10 sec trials. All correct 5 sec delay trials were used for training and all 10 sec trials for decoding. Only the delay time bins were used. In order to account for the difference in length between training and decoding delays, the decoder was applied on every second time bin of the 10 sec trials. Using this scheme, decoding will be efficient only if the initial delay sequence is expanded symmetrically in the extended delay to cover the doubled duration, so that a time-cell with a field at, say, 3 sec of the initial delay, will shift its field to 6 sec in the extended delay, and so on. If time-cells are randomly shuffled in the extended delay, this scheme is not expected to yield significant decoding accuracy at any time bin. Baseline was created by randomly shuffling the identities of the cells in the 10 sec delay trials, with 1000 repetitions and applying to decoder at each repetition.

For analysis across training days, Bayesian decoders were built as before, but only correct trials from the last 1/3 trials of each day were decoded. Therefore, as performance improved with training, more trials were used for training the decoder and for decoding each day. In naïve mice, the same process was followed and again only correct trials were decoded. Since these mice typically never licked for reward, correct trials refer to match-trials (where the mouse was not supposed to lick), implying that half of the last 1/3 trials were decoded. Baseline was created for each mouse each training day, following the same shuffling procedure as before. Mean baselines were computed by averaging across mice each day. For each day, significance over baseline was extracted by comparing pooled Bayesian data points from all mice with pooled chance values from each shuffling iteration in each mouse.

#### Support Vector Machine

A binary support vector machine (SVM) classifier was used to assess the odor stimulus in a trial based on the collective activity of all odor or time-cells. The same method as in Bayesian decoding was used to split trials into the training set and prediction set. All trials were split into two groups based on their odor identity. Again, only the correct trials were used from the training pool, whereas all trials from the decoding pool were decoded. Only sessions with at least 2 training trials of each kind were analyzed. The decoder was applied on the collective firing rates of all sequence cells, averaged either over the entire odor-delay interval, or the odor presentation timepoints or delay timepoints accordingly, or only over the firing field time point of each cell. The corresponding mean rate of each cell was z-scored over all trials. The classifier was trained using a radial basis function kernel with scale σ = 11 and box constrain parameter C = 15505. These values were acquired by Bayesian optimization of the SVM classifier over the two parameters, using the odor cell firing rates over odor time-bins from one imaging session and a 10-fold cross-validation partition for data. Odor-prediction accuracy refers to the percentage of correct odor-predictions over all predicted trials and is averaged over all imaging days analyzed. Chance baselines where computed by randomly shuffling the cell identities for each predicted trial with 1000 repetitions and applying the SVM classifier on the shuffled data for each repetition. The same classifier parameters and method were applied when using SVM on individual sequence-cells instead of collective sequence-cell groups (Figure 3E). Chance baseline for each cell was computed by shuffling the identity of both training and predicted trials with 50 repetitions and applying the SVM classifier on each repetition.

For cross-days analysis, we applied the SVM classifier on the firing rates of the recurring sequence-cells between each pair of days, following the same process as with Bayesian decoding (see above). The classifier was applied separately on recurring cells that were odor-cells on Day 1 and those that were time-cells, irrespective of their field on Day 2. Baseline was created by randomly shuffling the odor-identity of trials on Day 2, with 1000 repetitions and applying to decoder at each repetition. Decoded-odor accuracy by either cell group was pooled over all day pairs of a given distance and their mean value was computed and plotted (Figure 4J).

#### Statistical Analysis

Unless otherwise stated, most statistical tests between distribution averages were performed under the Wilcoxon median test (‘WT’) if the corresponding distributions were not sufficiently close to normality under the Lillieform normality test (p > 0.05). Otherwise, a paired-sample t-test was used. P-values were corrected for multiple comparisons wherever necessary. ‘FDR’ across the text refers to FDR-correction, and ‘Bonferroni’ to Bonferroni correction.

